# Tumor-induced endothelial RhoA activation mediates tumor cell transendothelial migration and metastasis

**DOI:** 10.1101/2024.09.22.614304

**Authors:** Md Sanaullah Sajib, Fatema Tuz Zahra, Margarita Lamprou, Racheal Grace Akwii, Jee Hyun Park, Manuel Osorio, Paul Tullar, Colleen L. Doci, Chonghe Zhang, Stephan Huveneers, Jaap D. Van Buul, Ming-Hai Wang, Maciej M. Markiewski, Sanjay K. Srivastava, Yi Zheng, J. Silvio Gutkind, Junhao Hu, Ulrich Bickel, Dean Y. Maeda, John A. Zebala, Michail S. Lionakis, Scott Trasti, Constantinos M. Mikelis

## Abstract

The endothelial barrier plays an active role in transendothelial tumor cell migration during metastasis, however, the endothelial regulatory elements of this step remain obscure. Here we show that endothelial RhoA activation is a determining factor during this process. Breast tumor cell-induced endothelial RhoA activation is the combined outcome of paracrine IL-8-dependent and cell-to-cell contact β_1_ integrin-mediated mechanisms, with elements of this pathway correlating with clinical data. Endothelial-specific RhoA blockade or *in vivo* deficiency inhibited the transendothelial migration and metastatic potential of human breast tumor and three murine syngeneic tumor cell lines, similar to the pharmacological blockade of the downstream RhoA pathway. These findings highlight endothelial RhoA as a potent, universal target in the tumor microenvironment for anti-metastatic treatment of solid tumors.

## Introduction

Metastasis, the dissemination of tumor cells to distant sites of the body, is the bottleneck of cancer research, and it is responsible for the majority of cancer-related deaths ^1,2^. The role of tumor microenvironment during metastasis is critical and presents an attractive therapeutic target due to its higher genetic stability compared to the tumor cells and the decreased susceptibility to therapeutic resistance mechanisms ^3^. During the metastatic process, the disseminating tumor cells have to re-adjust, pass and survive through several restrictive steps. These include the transendothelial migration for entrance in the tumor vasculature (intravasation), the adherence to the endothelium or entrapment in the capillaries at the metastasis-target organ and the transendothelial migration from the vasculature to the surrounding tissues (extravasation) ^1,4–6^. Therefore, the migration of tumor cells through the endothelial monolayer (intravasation or extravasation) is a determining step for the metastatic outcome. Here, we explore the role of endothelial RhoA in transendothelial tumor cell migration and metastatic outcome.

RhoA is one of the best-studied members of the Rho GTPase family, a highly conserved family among eukaryotes, also found in plants and fungi ^7^. It is an important mediator of actomyosin dynamics and participates in several fundamental cellular processes, ranging from cell migration, adhesion, polarization, proliferation to cell cycle progression and gene expression ^7–9^. RhoA is activated by extracellular factors binding to transmembrane receptors, such as G protein-coupled receptors (GPCRs), tyrosine kinase receptors and adhesion molecules ^10–12^. Similar to all GTPases, RhoA activation is balanced by the coordinated activity of guanine nucleotide exchange factors (GEFs), GTPase-activating proteins (GAPs) and guanine nucleotide dissociation inhibitors (GDIs). GEFs promote the GDP to GTP exchange, leading to an active GTPase state, GAPs enhance the intrinsic GTPase activity, thus inactivating the protein and GDIs sequester GTPases in the cytosol in a GDP-bound state, blocking spontaneous activation ^10,13^.

The specific functions regulated by RhoA are largely tissue-dependent. In the vasculature, RhoA controls important functions of the endothelial cells, including contraction, motility, proliferation and inflammation and its dysregulation is associated with vascular diseases ^14^. Endothelial permeability is an important vascular function mediated by RhoA. GPCR- and integrin-mediated endothelial RhoA activation induces endothelial permeability and vascular leakiness ^15–18^. The molecular mechanism involves the downstream activation of Rho kinase (ROCK) and myosin light chain (MLC) phosphorylation. MLC activation can occur both directly, through MLC kinase phosphorylation, or indirectly, through phosphorylation and subsequent inhibition of MLC phosphatase, promoting actomyosin contractility and stress fiber formation ^17,19–23^. RhoA-ROCK pathway-mediated endothelial hyperpermeability and vascular leakiness have been implicated in various pathological conditions ^24–27^ and we have previously shown that endothelial RhoA inhibition can potently block vascular leakiness and partially rescue from anaphylactic shock-related death ^17^. Despite the stimulating effect on endothelial permeability, RhoA activation has been shown to provide a protective role against endothelial permeability, such as acting downstream of Ang1 and blocking VEGF-induced permeability through a different downstream effector, mDia ^28^.

The regulatory role of endothelial RhoA in vascular permeability prompted us to explore its involvement in transendothelial migration and metastasis. We identified that the highly metastatic human breast tumor cells MDA-MB-231 and three syngeneic murine tumor cells (EO771 breast cancer cells, B16-F10 melanoma cells and Lewis Lung Carcinoma cells) activate endothelial RhoA. Tumor-induced endothelial RhoA activation seems to be the synergistic outcome of paracrine and cell-to-cell contact mechanisms. Secretome analysis from a panel of breast tumor cell lines correlated the transmigratory potential with the secretion levels of certain inflammatory mediators, among which, IL-8 was sufficient to drive p115-RhoGEF-mediated RhoA activation in the endothelial cells and endothelial permeability, enhancing tumor transendothelial migration. Tumor -endothelial cell-to-cell contact synergistically drove endothelial RhoA activation through an endothelial β_1_ integrin-mediated mechanism. Pharmacological blockade of endothelial RhoA and knockdown experiments confirmed the significance of endothelial RhoA in transendothelial migration of different solid tumors and endothelial-specific RhoA deficiency inhibited the metastatic potential in the syngeneic models. The downstream RhoA effector, Rho kinase or ROCK, is a known permeability mediator ^15,17^. Pharmacological inhibition of endothelial ROCK potently blocked transendothelial migration and the metastatic potential of murine syngeneic and human tumor cell lines, highlighting the endothelial RhoA pathway as an important target for anti-metastatic therapy.

## Materials and methods

### Cell Lines and Primary Cells

#### Human Umbilical Vein Endothelial Cells

Human Umbilical Vein Endothelial Cells (HUVECs) were isolated from collected umbilical cords from informed consent donors as per regulations following Institutional Review Board (IRB)-approved protocols A15-3891 from Texas Tech University Health Sciences Center Institutional Review Board and 13725 from Research Ethics Committee of the University of Patras, as previously described ^29^. HUVECs were cultured on gelatin-coated (1%) plates in M199 medium (Fisher Scientific, Cat# MT10060CV) supplemented with 15% Fetal Bovine Serum (FBS, Thermo Fisher Scientific, Cat# 10438026), 150 μg/ml Endothelial Cell Growth Supplement (ECGS, Fisher Scientific, Cat# CB-40006B), 5 U/ml heparin sodium (Hospira, Cat# NDC-63739-920-11) and 1X Antibiotic-Antimycotic solution (Fisher Scientific, Cat# 15240-062). All experiments were performed in HUVECs from at least three different donors and the cells were used between passages 1 and 6.

#### Mouse Lung Endothelial Cells

Mouse lung endothelial cells were isolated as described in the Method Details below. Mouse endothelial cells were cultured on 0.4% gelatin-coated dishes in M199 medium supplemented with 15% Fetal Bovine Serum (FBS), 150 μg/ml Endothelial Cell Growth Supplement (ECGS), 5 U/ml heparin sodium and 1X Antibiotic-Antimycotic solution.

#### EAhy926 Cells

EAhy926 cells were purchased from ATCC and cultured in DMEM (Fisher Scientific, Cat# 11-995-073) supplemented with 10% Fetal Bovine Serum (FBS) and 1X Antibiotic-Antimycotic solution, as previously described ^17^.

#### MDA-MB-231 Cells

MDA-MB-231 GFP+ and MDA-MB-231 Luciferase-2A-RFP+ cells were purchased from Cell Biolabs and GenTarget, respectively. MDA-MB-231 GFP+ were cultured in DMEM supplemented with 10% Fetal Bovine Serum (FBS) and 1X Antibiotic-Antimycotic solution according to supplier’s instructions. MDA-MB-231 Luciferase-2A-RFP+ cells were cultured in L-15 medium (Himedia Labs, Cat# AL011S), supplemented with 10% FBS, 0.1 mM non-essential amino acid solution (Thermo Fisher Scientific, Cat# 11140-050), 2 mM Glutamine (Thermo Fisher, Cat# 25030-081) and 1X Antibiotic-Antimycotic solution, according to supplier’s instructions.

#### MDA-MB-468 Cells

MDA-MB-468 cells were purchased from ATCC and cultured in DMEM supplemented with 10% Fetal Bovine Serum (FBS) and 1X Antibiotic-Antimycotic solution.

#### T-47D Cells

T-47D cells were purchased from ATCC and cultured in RPMI-1640 (Fisher Scientific, Cat# 11-875-119) supplemented with 10% Fetal Bovine Serum (FBS), 1X HEPES (Thermo Fisher Scientific, Cat# 15630080) and 1X Antibiotic-Antimycotic solution.

#### SUM52PE Cells

SUM52PE cells were obtained from the MH Wang lab and cultured in DMEM supplemented with 10% Fetal Bovine Serum (FBS) and 1X Antibiotic-Antimycotic solution.

#### EO771 Cells

EO771 cells were obtained from the M Markiewski lab and cultured in RPMI-1640 supplemented with 10% Fetal Bovine Serum (FBS), 1X HEPES and 1X Antibiotic-Antimycotic solution.

#### B16-F10 Cells

B16-F10 cells were obtained from ATCC and cultured in DMEM supplemented with 10% Fetal Bovine Serum (FBS) and 1X Antibiotic-Antimycotic solution.

#### Lewis Lung Carcinoma Cells (LLC)

LLC cells were obtained from ATCC and cultured in DMEM supplemented with 10% Fetal Bovine Serum (FBS) and 1X Antibiotic-Antimycotic solution.

#### BEAS-2B Cells

Human lung epithelial cell line BEAS-2B was obtained from ATCC and cultured in MEM complete medium containing 10% FBS and 1X Antibiotic-Antimycotic solution.

All tumor cells were authenticated with short tandem repeat (STR) assay from Texas Cancer Cell Repository (Texas Tech University Health Sciences Center) and ATCC. All cell cultures were maintained at 37°C and 5% CO_2_ and were tested for mycoplasma.

### Mice

All animal studies were performed according to the approved protocols of Institutional Animal Care and Use Committee (IACUC) of Texas Tech University Health Sciences Center (TTUHSC) and of the Directorate of Veterinary Medicine of the Region of Western Greece. For the experimental metastasis model with the human MDA-MB-231 cells NOD SCID mice were used, whereas for all the Fasudil experiments we used wild-type mice of the C57BL/6J background. For the syngeneic EO771 breast tumor cell line only females were used, whereas for the B16-F10 and the LLC cell lines both male and female mice were used. All mice were 8 weeks of age or older. The generation of the *RhoA* floxed mice was previously reported ^30^. These mice were crossed with mice carrying the *Tomato-GFP* reporter and the endothelial-specific RhoA deficiency was obtained by crossing them with mice carrying a tamoxifen-inducible Cre-mediated recombination system driven by the *Cdh5* promoter (*Cdh5-CreERT*^2^), as previously reported ^17^. To induce Cre-mediated gene deletion, Tamoxifen dissolved in Miglyol was intraperitoneally (IP) injected for five consecutive days (2 mg Tamoxifen in 100 µl Miglyol/mouse/day). All littermate mice were treated and the ones without the promoter were used as control.

## Method Details

### Mouse lung endothelial cell isolation

Mouse lung endothelial cell isolation was performed as previously described ^17,29^. The isolation and the experiments took place after at least one week after the end of tamoxifen treatment. Lungs were removed, washed in DMEM with 10% FBS, minced under sterile conditions into pieces of size 1-2 mm^3^ and followed by digestion with Collagenase Type I (2mg/ml, Fisher Scientific, Cat# 17-100-017) for 2h at 37°C with occasional agitation. The cellular digest was filtered through a 70 μm cell strainer, centrifuged at 450 g and the cells were plated on gelatin-coated dishes with aforementioned media (M199 medium supplemented with 15% Fetal Bovine Serum (FBS), 150 μg/ml Endothelial Cell Growth Supplement (ECGS), 5 U/ml heparin sodium and 1X Antibiotic-Antimycotic solution) (day 0). On days 1 to 5 the plates were washed twice with PBS (Fisher Scientific, Cat# SH30256FS) and the medium was replaced daily. On day 5 the first endothelial purification took place with 10 min incubation with pre-coated magnetic beads with rat anti-mouse ICAM-2 mAb (3C4, BD Biosciences, Cat# 553325) at room temperature under continuous agitation, followed by two PBS washes and trypsinization. The bead-bound cells were recovered with a magnet, washed four times, resuspended in endothelial growth media and plated on gelatin-coated dishes. After the second purification (on day 10, depending on the confluency) the cells were used for analysis.

### *In-vitro* transendothelial migration model

Transwell inserts with 6.5 mm diameter and 8 µm pore-sized polycarbonate membrane (Fisher Scientific, Cat# 07-200-150) were coated with 400 µl Collagen type I (in 0.2 N acetic acid, Fisher Scientific, Cat# CB354249) for 20 min at 37°C. After incubation, collagen was removed and the inserts were washed with PBS to remove excess collagen. The wells of 24-well plate were filled with 600 µl medium and 2×10^5^ EAhy926 cells or HUVECs, suspended in 100 µl medium, were added into the insert. The cells were allowed to grow for 4 to 6 days, depending on the experiment, with medium replacement every other day. Cells were treated with relevant antibodies (15 min) or pharmacological agents for the time indicated in figure legends. On the day of the experiment, the inserts were transferred to new wells with fresh medium. Tumor cells were either expressing fluorescence by lentiviral-mediated GFP/RFP expression or by labeling with CellTracker^TM^ Green CMFDA (Thermo Fisher Scientific, Cat# C7025) dye. In all experiments, 2×10^5^ tumor cells were added on the inserts and allowed to migrate through the endothelial layer for 8 to 24 h. At the end of the incubation period, the cells were fixed in 4% PFA (Electronic Microscopy Systems, Cat# 15714-S) for 10 min and stained with Hoechst (Thermo Fisher Scientific, Cat# H3570). The non-migrated cells on the dorsal side of the insert were removed with cotton swabs and membranes were separated from the inserts and mounted on glass slides. Approximately 7-10 images were taken per insert and the transmigrated tumor cells (with green or red fluorescence) were quantified. All experiments were performed at least three times unless stated otherwise.

### Transendothelial electrical resistance (TEER) experiments

Transendothelial impedance was measured with cellZscope (nanoAnalytics GmbH, Germany), according to the manufacturer’s instructions. Briefly, transwell inserts with 6.5 mm diameter and 3 or 8 µm pore-sized polycarbonate membrane were coated with 400 µl Collagen type I (in 0.02 N acetic acid) for 20 min at 37°C. After incubation, collagen was removed and the inserts were washed with PBS to remove excess collagen. The wells of 24-well plate were filled with 600 µl medium and 2×10^5^ EAhy926 cells were added to the transwells and cultured for 6 days, with medium change every other day. Transwells were transferred to the cellZscope instrument with 810 μl medium in the bottom well and 260 μl in the top one. After acquiring the baseline resistance reading of each well, either control medium, conditioned medium or cells (2×10^5^ EAhy926 or MDA-MB-231) were added and the measurement of resistance was continued for 8 h more.

### Cytokine Analysis

For the detection of a panel of 36 proinflammatory cytokines, the Bio-Plex Pro™ Human Inflammation Panel 1 immunoassay kit (Bio-Rad, Cat#171AL001M) was used and the cytokine array profiling was performed with a Biorad BioPlex 200 system using the Bioplex Manager Software package, according to manufacturer’s instructions. Briefly, breast tumor cell lines MDA-MB-231, MDA-MB-468, T-47D and SUM52PE were seeded in 100 mm^3^ plates and starved with 7 ml serum-free M199 medium. After 24 h, the supernatant was collected and centrifuged to get rid of cell debris. The collected conditioned media were stored at -80°C until analysis.

### RhoA pull-down assay

Endothelial RhoA activation was measured by RhoA pull-down experiments as previously described ^29^. The GST-RBD plasmid was a gift from Martin Schwartz (Addgene plasmid # 15247)^31^. For the time course experiments of the effect of tumor cell supernatant on endothelial RhoA activation 10 cm dishes with tumor cells in confluency were starved with 7 ml of M199 medium overnight. For the comparative study of endothelial RhoA activation from supernatant from different breast tumor cell lines, 2×10^6^ cells were cultured overnight in 10 cm dishes in full medium, the medium was replaced with 7 ml serum-free M199 medium the next day, and it was collected after 24 h of incubation. The supernatant of the conditioned medium after a 5 min centrifugation at 200 g was collected and processed for the endothelial incubation. For the effect of recombinant cytokines on endothelial RhoA activation, HUVECs were incubated with the identified, from the secretome analysis, concentration of each cytokine in M199 medium. For all experiments, HUVECs were plated in gelatin-coated 6-well plates and cultured up to 85-90% confluency. After 3 h of starvation, 1 ml of the conditioned medium or the recombinant cytokines in the appropriate concentration was added in the wells for the specific time points. After a quick wash with ice-cold PBS, HUVECs were lysed in RhoA kinase lysis buffer containing 20 mM HEPES, pH 7.4, 100 mM NaCl, 1% Triton X-100, 20 mM MgCl_2_, 10 mM EGTA, 40 mM β-glycerol phosphate, 1 mM phenylmethylsulfonyl fluoride (PMSF), 10 µg ml^-1^ Aprotinin, 10 µg ml^-1^ Leupeptin, 1 mM Na_3_VO_4_ and 1 mM dithiothreitol (DTT). Cell lysates were centrifuged at 13,200 g for 5 min at 4 °C, and the supernatants were incubated with 30 μl of previously conjugated with glutathione beads GST-RBD fusion protein, at 4 °C for 30 min and washed with lysis buffer. The beads, as well as 50 µl of the supernatant from each sample, were mixed with 2X SDS-polyacrylamide gel electrophoresis sample buffer and analyzed by western blot analysis for active and total RhoA respectively, as described below.

### GEF pull-down assay

GEF pull down was performed as previously described ^32^ with some modifications. The pGEX-4T1-RhoA G17A plasmid was a gift from Rafael Garcia-Mata (Addgene plasmid # 69357) ^32^. HUVECs were starved with M199 for 4 h before treating the cells with 10 ng/ml of recombinant IL-8 for 3 min. Cells were lysed in RhoA kinase lysis buffer (described above). Cell lysates were centrifuged at 13,200 g for 5 min at 4 °C, and the supernatants were incubated with 30 μl of previously conjugated with glutathione beads GST RhoA^G17A^ fusion protein, at 4°C for 30 min and washed with lysis buffer. The beads, as well as 50 µl of the supernatant from each sample, were mixed with 2X SDS-polyacrylamide gel electrophoresis sample buffer and analyzed by western blot analysis for active and total RhoA GEFs respectively, as described below.

### Immunoblot analysis

Immunoblot analysis was performed as previously described ^33^. The cells were analyzed in RIPA buffer (10 mM Tris-HCl, 140 mM NaCl, 1 mM EDTA, 0.5 mM EGTA, 0.1% sodium deoxycholate, 0.1% SDS and 1% Triton X-100), supplemented with protease and phosphatase inhibitor cocktail (Halt Protease and Phosphatase Inhibitor Cocktail). Cell lysates were centrifuged at 13,000 g for 10 min at 4°C and after protein quantification were mixed with 2X SDS-polyacrylamide gel electrophoresis sample buffer. To analyze IL-8 levels in transfected MDA-MB-231 cells, the secreted proteins in the cell culture medium were precipitated using trichloroacetic acid, as previously described ^34^: Briefly, cell supernatant was mixed with an equal amount of 20% Trichloroacetic acid. After 15-30 min incubation on ice, the sample was centrifuged at 10,000 g for 5 min at 4°C, the supernatant was removed and the pellet was washed in cold acetone and re-centrifuged. The process was repeated twice and the dry pellet was resuspended in SDS-polyacrylamide gel electrophoresis sample buffer. Samples were heated to 100°C for 5 min and centrifuged. Equal amounts of proteins were subjected to SDS-PAGE and transferred onto an Immobilon P, polyvinylidene difluoride membrane.

Primary antibodies were used in the following dilutions: β-actin (Cell Signaling Technology, Cat# 3700, 1:2,000), β-tubulin (Cell Signaling Technology, Cat# 2146, 1:2,000), RhoA (Cell Signaling Technology, Cat# 2117, 1:1000), Integrin β1 (Cell Signaling Technology, Cat# 34971, 1:1,000), Integrin β3 (Cell Signaling Technology, Cat# 4702, 1:1,000), GEF-H1 (Cell Signaling Technology, Cat# 4076, 1:1,000), p115-RhoGEF (Cell Signaling Technology, Cat# 3669, 1:1,000), PDZ-RhoGEF (Abcam, Cat# ab110059, 1:500), LARG (Abcam, Cat# ab136072, 1:500), IL-8 (Thermo Fisher Scientific, Cat# 701232, 1:200). Secondary antibodies were horseradish peroxidase-conjugated anti-rabbit (Southern Biotech, Cat# 4010-05) and anti-mouse (Southern Biotech, Cat# 1010-05) IgGs (1:50,000 for both). Activation was assessed by the ratio of active versus total RhoA or GEFs for the pull-down experiments and expression by the ratio of protein versus β-actin or tubulin. Densitometric quantification of scanned western blots was performed using ImageJ analysis software (National Institutes of Health).

### Boyden Chamber Cell Migration Assay

Cell migration was performed as previously reported ^35^, using a 48-well Boyden chamber with an 8-μm pore size polyvinyl pyrrolidone-free polycarbonate membrane (NeuroProbe) coated with collagen. Tumor cells were added to the upper chamber 48 h post-transfection, and DMEM with 10% FBS was added to the lower chamber. After incubation for 6 h at 37°C, the cells on the upper surface of the membrane were removed, and the cells at the lower surface were fixed with methanol and stained with hematoxylin. The cells were counted under a bright-field microscope (Microscoptics, IV-900).

### siRNA Transfection

The following siRNAs were used: RhoA #1, RhoA #2, IL-8, p115-RhoGEF, Integrin β1 and Integrin β3 (Thermo Fisher Scientific, Cat#s758, Cat#s759, Cat#s7328, Cat#119421, Cat#s7575 and Cat#112581 respectively) or a negative control (Cat#4390846). Cells were transfected with siRNAs with DharmaFECT1 (Dharmacon) transfection reagent at 50 nM concentrations, according to manufacturer’s instructions, with some modifications. Briefly, cells were seeded in full media in 6-well plates and grown up to 90% confluency. For HUVECs media were replaced to antibiotic- and serum-free starvation media 1 h prior to transfection, whereas all cell lines (EAhy926, MDA-MB-231) were starved with antibiotic- and serum-free media overnight and replaced to fresh antibiotic- and serum-free media 1 h prior to transfection. 6 h after addition of the transfection complex medium was replaced to full medium and 48 h post-transfection, the cells were either lysed for western blot analysis or were used for other experiments as indicated in figure legends. For trans-endothelial migration experiments, cells were trypsinized and seeded in transwells 24 h post-transfection.

### Immunofluorescence

For endothelial-tumor cell co-culture experiments HUVECs were cultured on sterile glass slides coated with 1% gelatin 24 h prior to seeding of the GFP-expressing MDA-MB-231 tumor cells. 24 h later, the cells were fixed with 4% PFA for 10 min at room temperature (RT), permeabilized with 0.2 % Triton X-100 for 10 min at RT and blocked with PBS containing 3% BSA for 1 h at RT. The cells were immunostained for anti-GTP-RhoA (NewEast Biosciences, Cat# 26904, 1:100), anti-integrin β1 (Abcam, Cat# ab30394, 1:100) and pMLC (Cell Signaling Technologies, Cat# 3671, 1:50) overnight at 4°C with the primary antibodies diluted in PBS containing 3% BSA. Then the cells were immunostained with the appropriate secondary antibodies (Thermo Fisher Scientific, Donkey anti-mouse Alexa Flour 594 Cat# A-21203; Donkey antimouse Alexa Flour 647 Cat# A-31571; Donkey anti-rabbit Alexa Flour 647 Cat# A-31573, 1:500), diluted in PBS containing 3% BSA, for 2 h at RT. To visualize F-actin, the cells were stained with Alexa Flour 594 Phalloidin overnight at 4°C. Nuclei were stained with Hoechst (Life Technologies, Cat# H3570, 1:1000 in PBS). Fluorescence images were recorded with a multiphoton microscope (A1R; Nikon, NY, USA) in the confocal mode with Plan-Apochromat Lamda 20X/0.75 numerical aperture and Plan Apochromat IR 60X/1.27 numerical aperture water immersion objective lenses. The microscope and image acquisition were controlled by Nikon NIS software.

### Intravital imaging of mouse ear skin

One day before the experiment, the mouse was anesthetized by isoflurane inhalation using the LARC-A E-Z Anesthesia induction chamber (0.5-2% isoflurane, flow=1L/min; Euthanex Corp, Palmer, PA) and the ear hair was removed from the imaging area by gently applying adhesive masking tape a few times. The day of the experiment, the mouse was anesthetized by isoflurane inhalation and the mouse ear was stabilized on a customized ear-holder platform with adhesive masking tape in the periphery of the ear. GFP was excited with a 488-nm laser, whereas m-Tomato was excited with a 561-nm laser. Intravital microscopy was acquired through an with a multiphoton microscope (A1R; Nikon, NY, USA) in the confocal mode with Plan-Apochromat Lamda 20X/0.75 numerical aperture air objective lens. The microscope and image acquisition were controlled by Nikon NIS software.

### Real Time RT-PCR Analysis

Total RNA was isolated from siRNA transfected MDA-MB-231 at 48 h post-transfection using Direct-zol RNA Miniprep Plus kit (Zymo Research, Cat# R2071), according to manufacturer’s instructions. 1 µg of RNA was used for synthesis of cDNA using the Verso cDNA synthesis kit (Thermo Fisher). Real-time RT-PCR was performed using 1 µl of cDNA for the gene-specific primers for IL-8 and β-actin using a standard SYBR green Taqman protocol (Thermo Fisher Scientific, Cat# 4309155) and Bio-Rad CFX96 Real-Time PCR machine. Fold-expression levels of IL-8 were normalized with β-actin and expressed as ΔΔCT values from three replicates from each experiment.

### ELISA

IL-8 levels in MDA-MB-231 tissue culture serum-free conditioned medium 48 h after siRNA transfection were identified with the IL-8 Human ELISA kit (Thermo Fisher Scientific, Cat# KAC1301) according to manufacturer’s instructions.

### Experimental Metastasis Model

For experimental metastasis studies, NOD SCID mice (Jackson Laboratory, Cat# 001303) were used for human MDA-MB-231 breast tumor cells and C57BL/6J mice (Jackson Laboratory, Cat# 000664) were used for EO771, B16-F10 and LLC murine syngeneic tumor cells. The endothelial-specific, conditionally RhoA deficient mice were in the C57BL/6 background as well. For breast tumor cells (MDA-MB-231 and EO771), only female mice were used, whereas male and female mice were used for B16-F10 and LLC cells. All mice were aged 8 weeks or older.

For both intravenous and intracardiac experimental metastasis models, 1×10^5^ tumor cells in 100 µl sterile PBS were administered. For intravenous metastasis model, the tumor cells were injected in the lateral caudal vein, and for the intracardiac model, they were injected in the left ventricle of the heart with the help of a stereotactic instrument. Injection accuracy was evaluated by a pulsatory flash of bright-red blood into the syringe upon little retraction of the plunger prior to injection. In experiments evaluating Fasudil’s effect on metastatic potential, PBS versus 20 mg/kg Fasudil in 100 μl sterile PBS were administered intraperitoneally (IP) daily (starting the day before the inoculation), for the duration of the experiment. SX-682 was obtained under a Cooperative Research and Development Agreement with Syntrix Pharmaceuticals. Control chow and chow formulated with SX-682 (drug added to expose mice to 500 mg/kg body weight/day) was obtained from Research Diets, Inc and were administered via oral gavage.

For metastatic progression evaluation, 3 mg of luciferin potassium in 100 µl PBS were administered intraperitoneally (IP), and animals were imaged by IVIS Lumina XR (Caliper Life Sciences, Inc.). Upon completion of the experiment, mice were humanely euthanized, lungs were dissected and the number of metastases was identified prior to histological analysis.

### Spontaneous Metastasis Model

For spontaneous metastasis studies, endothelial-specific conditional RhoA-deficient female mice aged 7 weeks or older were used. Puromycin-resistant 1×10^6^ EO771-luc cells in 100 μl sterile PBS were administered in the 3rd mammary gland of the female mice. Primary tumors were aseptically removed when they reached to 10 mm of diameter and sutured. To evaluate metastatic progression, 3 mg of luciferin potassium in 100 μl PBS was administered intraperitoneally (i.p.) and animals were imaged by IVIS Lumina XR (Caliper Life Sciences, Inc.). On day 50 after primary tumor inoculation, mice were humanely euthanized, lungs were isolated, and number of metastases was quantified. Also, 0.5 ml of blood was collected in heparin-coated tubes and the circulating tumor cells (CTCs) were isolated using previously described protocol^36^. CTCs were selected using puromycin (4 μg/ml) and images were obtained by IVIS using 0.1667 mg/ml of luciferin in PBS solution.

### Colony formation assay for extravasation

To detect extravasation efficiency, mice were intraperitoneally (i.p.) pre-treated with fasudil (20 mg/kg). 2 hours after fasudil treatment, puromycin-resistant 1×10^6^ EO771-luc breast cancer cells were intravenously injected in the tail vein. Lungs were harvested after 24 hour and single cell suspension was prepared. Cells were cultured in 37°C and 5% CO_2_ overnight in RPMI-1640 supplemented with 10% Fetal Bovine Serum (FBS) and 1X Antibiotic-Antimycotic solution. The following day the cells were treated with puromycin (4 μg/ml) to select the tumor cells. Medium was changed every 3rd day till the surviving cells formed colonies. When the colonies were sufficiently large, they were washed with PBS, fixed with 10% formalin for 15-30 min, and stained with 0.01% (w/v) crystal violet in water for 30-60 min, as previously described^37^ . The excess stain was removed with water and the plates were air-dried at room temperature. Colonies containing more than 50 individual cells were quantified with ImageJ.

### Histopathology and Immunohistochemistry

Routine histological analysis took place after lung dissection and IVIS imaging. At necropsy, lungs were inflated with 10% neutral buffered formalin (NBF, Millipore Sigma, Cat# HT501128) to preserve the normal architecture and then immersion fixed in 10% NBF for 24 h. Tissues were processed routinely for histology and embedded in paraffin. Tissue sections were cut on a rotary microtome at 5 microns, mounted on slides, stained with Hematoxylin & Eosin, and coverslipped. Photomicrography was obtained with a Nikon Eclipse Ni using NIS Elements software.

### Data mining

Kaplan–Meier plotter ^38^ (http://www.kmplot.com/analysis/) was used to determine the correlation of *IL-8*, *MDNCF*, *TNF-R1* and *BAFF* gene expression with distant metastasis-free survival of breast tumor patients. The parameters used to generate the graphs are listed in the supplemental text.

### Statistics

Statistics were performed with the GraphPad Prism 8 software. Unpaired two-tailed t-test with Welch’s correction was used to compare the means of two independent groups, and one-way ANOVA followed by Dunnett’s multiple comparisons test was used to compare the means of more than two independent groups of normally distributed data.

## Results

### Tumor cell-derived paracrine mediators induce endothelial permeability and activate endothelial RhoA

While the mechanisms of transendothelial migration have not been adequately elucidated ^39–41^, induction of vascular permeability has been established as a determining factor for the metastatic outcome ^41,42^. To assess whether tumor cell-derived paracrine mediators induce vascular permeability, endothelial cells were cultured in transwells (Figure 1A), and transendothelial electrical resistance (TEER) was measured in the presence of serum-free fresh or conditioned medium from tumor or endothelial cells ^43^. Addition of medium increased the TEER values in all groups, however addition of the tumor cell-(MDA-MB-231)-derived conditioned medium presented lower TEER values than of the control (M199) and the endothelial cell-(EAhy926)-derived conditioned medium (Figure 1B). Similar results were obtained when the TEER values were measured in a transwell-based co-culture model of transendothelial migration, where the tumor cells can transmigrate through an endothelial monolayer (Suppl. Figure 1A). Addition of endothelial cells (EAhy926) on the existing endothelial monolayer (dorsal part of the membrane) increased the TEER value, thus decreasing permeability (Suppl. Figure 1B). However, addition of metastatic breast tumor cells (MDA-MB-231) induced endothelial permeability, demonstrated by TEER value decrease, significantly higher than the control levels, suggesting that tumor cells or their secretome can breach the endothelial monolayer integrity (Suppl. Figure 1B). As endothelial RhoA is a known inducer of vascular permeability ^19,24,25^, we explored the effect of MDA-MB-231 secretome on endothelial RhoA activation by treating primary human umbilical vein endothelial cells (HUVEC) with serum-deprived conditioned medium from MDA-MB-231 tumor cells. RhoA pull-down experiments demonstrated a significant increase in RhoA activation of HUVECs (Figures 1C,D). To evaluate whether tumor cell-induced paracrine effect on endothelial RhoA activation correlates with transmigration and metastatic potentials, we selected four human breast tumor cell lines with reported differences in metastatic potential ^44^ and tested their transmigration efficiency in the transmigration *in vitro* model. The MDA-MB-231 cells migrated the most and the T-47D the least, while the SUM52PE did not migrate in this model (Figures 1E,F). Conditioned media from these cell lines showed a comparable, to the migrating potential, trend of endothelial RhoA activation (Figures 1G,H). Tumor cells secrete a large number of vasoactive compounds that are known to increase vascular permeability ^3,45^ and our RhoA activation data denoted potential qualitative and quantitative differences in the secretome of these cell lines. To investigate candidates responsible for the paracrine endothelial RhoA activation, we explored the secretion levels of known inflammatory mediators from each of these cell lines (Figure 1I and Suppl. Figure 1C). The analysis revealed that the expression of interleukin 8 (IL-8), B-cell activating factor (BAFF) and soluble tumor necrosis factor receptor 1 (sTNF-R1) was highest in the MDA-MB-231 cells, consistent with their transmigration potential.

**Figure 1.**
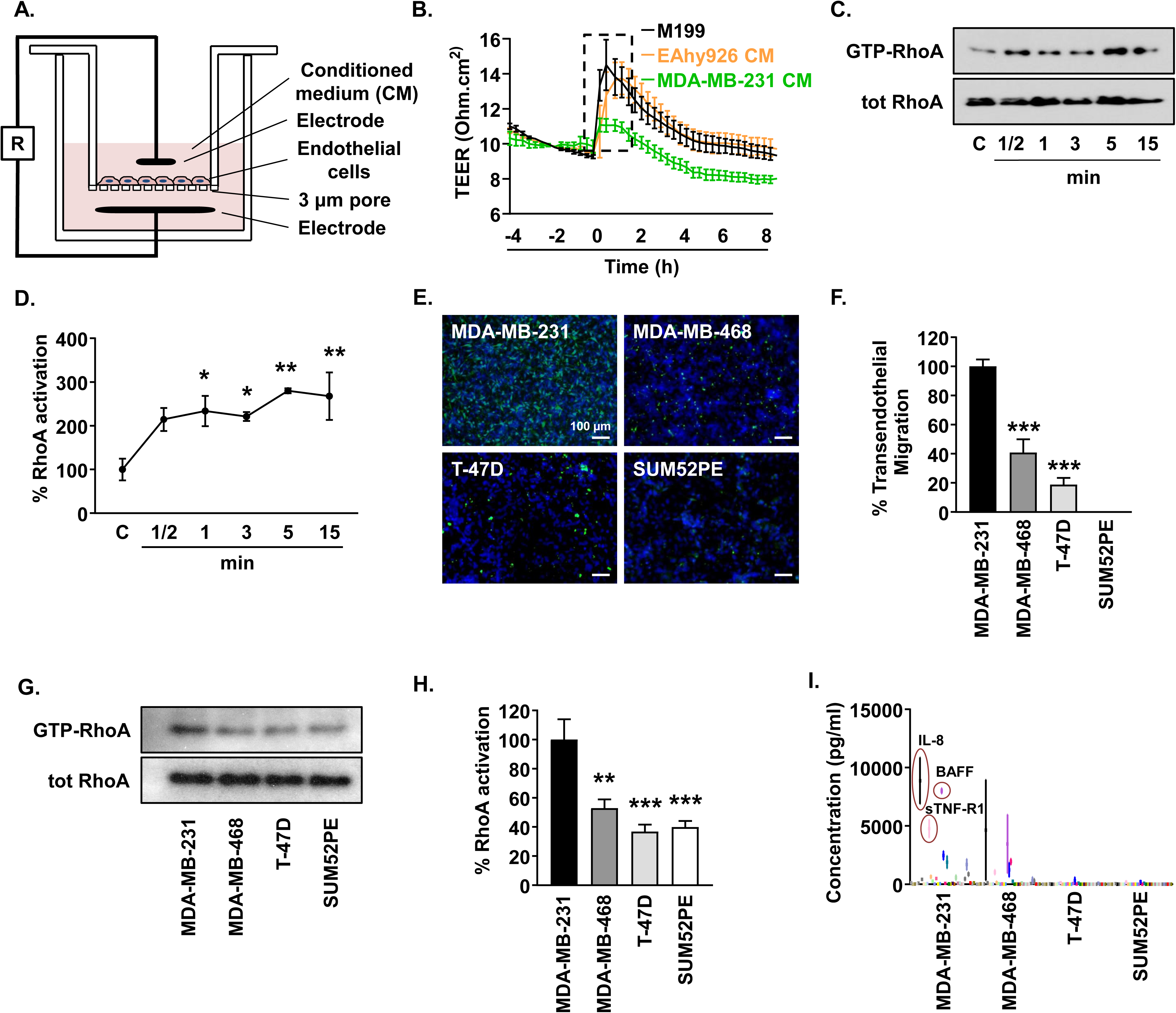
Tumor cell-derived paracrine mediators induce endothelial permeability and activate endothelial RhoA. (A) Schematic diagram of *in vitro* transendothelial electrical resistance (TEER) setting and measurement. (B) Measurement of TEER across endothelial monolayer before and after the addition of M199-serum-free condition medium (CM) from endothelial (EAhy926) or cancer (MDA-MB-231) cells or Μ199 medium (control). Dotted box designates the increase of TEER following conditioned or control medium addition. n = 2. (C and D) Representative images (C) and quantification (D) of RhoA activation in HUVECs upon incubation with serum-free M199 or conditioned medium from MDA-MB-231 cancer cells at various time points; n = 9. (E and F) Representative images (E) and quantification (F) of trans-endothelial migration of GFP expressing or green fluorescent dye-labeled human breast cancer cells through an endothelial (EAhy926) monolayer on collagen-coated transwell inserts after 24 h (nuclei = blue); n = 2. Scale bars, 100 µm. (G and H) Representative western blot images (G) and quantification (H) of RhoA activation in HUVECs upon incubation with serum-free conditioned medium from MDA-MB-231, MDA-MB-468, T-47D and SUM52PE cancer cells; n = 3. (I) Multiplex analysis of secretion of human inflammatory cytokines in serum-free conditioned media from the aforementioned cancer cell lines; n = 3. Data in (B), (D), (F), (H) and (I) represent mean ± SEM. Data in (D), (F) and (H) were analyzed by one-way ANOVA. *P < 0.05; **P < 0.01; ***P < 0.001. See also Figure S1.

### Tumor cell-derived Interleukin-8 activates endothelial RhoA and facilitates tumor cell transendothelial migration

To identify whether IL-8, BAFF and sTNF-R1 are responsible for endothelial RhoA activation, we performed RhoA pulldown experiments on HUVECs using recombinant proteins at the same concentrations as secreted by the MDA-MB-231 tumor cells. sTNF-R1 and BAFF did not activate RhoA in HUVECs at the tested concentrations (Suppl. Figures 2A,B), contrary to IL-8, which caused significant RhoA activation at 3 and 5 min (Figures 2A,B). To explore the impact of IL-8 in tumor cell transendothelial migration, we performed a gain-of-function experiment, by adding recombinant IL-8 to the poorly migrating T-47D tumor cells. IL-8 (10 ng/ml) presence increased the transmigration efficiency of the T-47D cells (Figures 2C,D). This increase is not due to the effect of IL-8 on T-47D intrinsic migration, as motility of T-47D has been reported unaffected by IL-8 up to 100 ng/ml concentration^46^. Similarly, the instrinsic migration of SUM52PE was not increased by the presence of IL-8 in the boyden chamber-based cell migration model (Suppl. Figure 2C, DMSO group), but the transendothelial migration was significantly increased by IL-8 presence (Suppl. Figures 2D,E), suggesting that the increased migration is due to IL-8’s effect on the endothelial cells.

**Figure 2.**
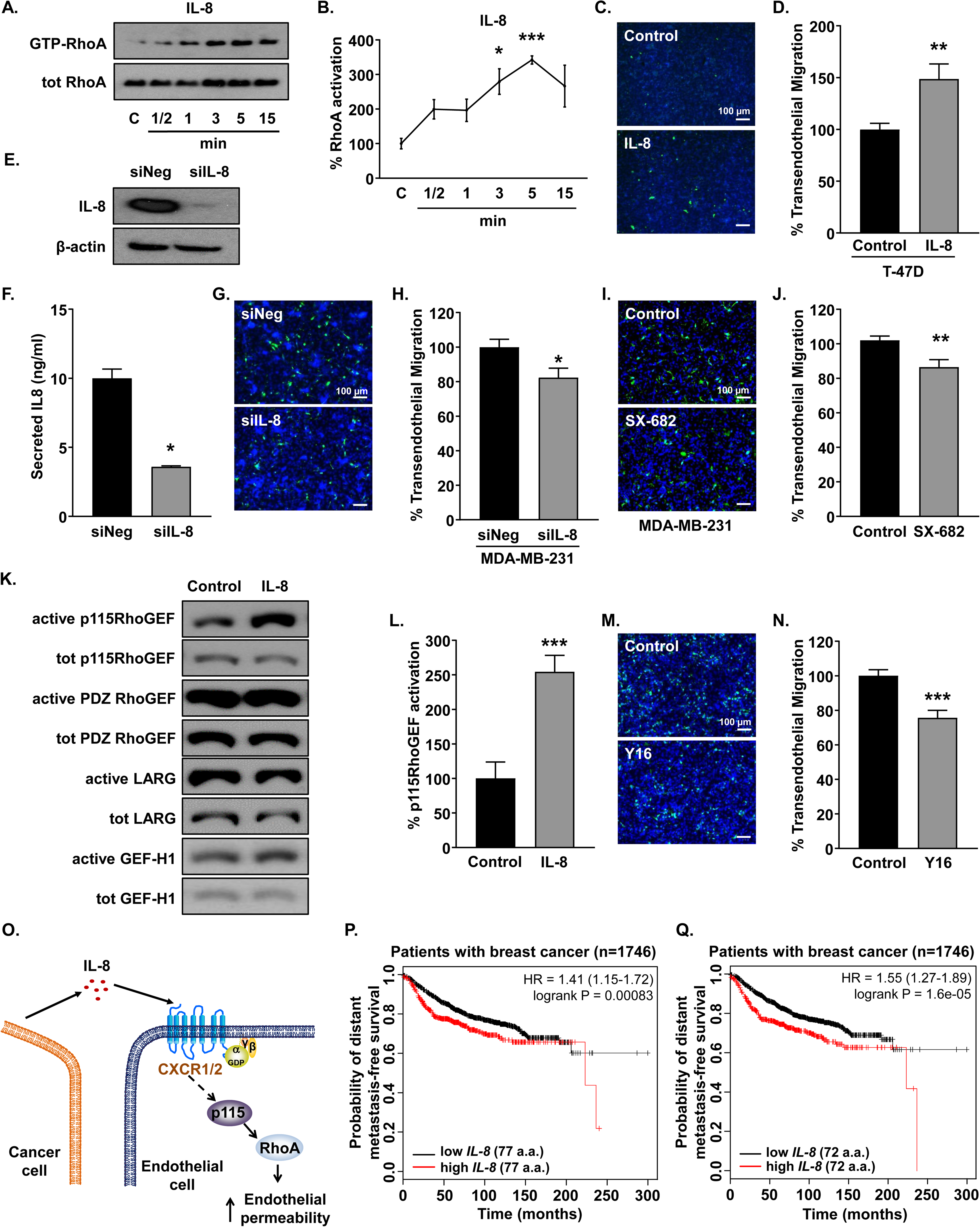
Tumor cell-derived interleukin-8 (IL-8) activates endothelial RhoA and facilitates cancer cell trans-endothelial migration. (A and B) Representative images (A) and quantification (B) of RhoA activation in HUVECs upon treatment of IL-8 (10 ng/ml), at different time points; n = 6. (C and D) Representative images (C) and quantification (D) of transendothelial migration of green fluorescent dye-labeled T-47D cancer cells through an EAhy926 monolayer on collagen-coated transwell inserts in absence or presence of recombinant IL-8 (10 ng/ml) after 24 h (nuclei = blue); n = 5. Scale bars, 100 µm. (E) Representative images of IL-8 knockdown in MDA-MB-231 by siRNA treatment (50 nM) at 48 h post-transfection; n = 2. (F) Quantification of secreted IL-8 in serum-free conditioned medium of siNeg- and siIL-8-transfected MDA-MB-231 at 48 h post-transfection, n = 2. (G and H) Representative images (G) and quantification (H) of trans-endothelial migration of GFP-expressing siNeg- and siIL-8-transfected MDA-MB-231 cancer cells through an EAhy926 monolayer on collagen-coated transwell inserts after 8 h (nuclei = blue); n = 3; Scale bars, 100 µm. (I and J) Representative images (I) and quantification (J) of trans-endothelial migration of GFP-expressing MDA-MB-231 cancer cells through an EAhy926 monolayer on collagen-coated transwell inserts without and upon treatment with the SX-682 inhibitor and 8h after the addition of the cancer cells (nuclei = blue); n = 4; Scale bars, 100 μm. (K) Representative images of IL-8-induced RhoGEF activation in HUVECs upon 5 min of IL-8 (10 ng/ml) incubation. (L) Quantification of IL-8-induced p115RhoGEF activation in HUVECs upon 5 min of IL-8 (10 ng/ml) incubation; n = 7. See also supplemental Figure 2E. (M and N) Representative images (K) and quantification (L) of transendothelial migration of GFP-expressing MDA-MB-231 tumor cells through an EAhy926 monolayer treated with vehicle or GhoGEF inhibitor Y16 (20 μM. n = 2; Scale bars, 100 μm. (O) Schematic diagram illustrating potential mechanism of endothelial RhoA activation by tumor-cell-derived IL-8. (P and Q) Metastasis-free survival of breast tumor patients, stratified on IL-8 expression. Two variants of IL-8, endothelial cell derived containing 77 amino acids (N) and monocyte-derived containing 72 amino acids (O) are shown. Data in (B), (D), (F), (H), (J), (L) and (N) represent mean ± SEM. Data in (B) were analyzed by one-way ANOVA and data in (D), (F), (H), (J), (L) and (N) were analyzed by Student’s unpaired t-test. Data in (N) and (O) was analyzed by Kaplan-Meier estimator. *P < 0.05; **P < 0.01; ***P < 0.001. See also Figure S2.

For a loss-of-function evaluation of IL-8 role, we knocked down IL-8 expression in the aggressive MDA-MB-231 tumor cells (Figures 2E-F and Suppl. Figures 2F,G). IL-8 knockdown did not affect the migration of MDA-MB-231 cells (Suppl. Figures 2H,I), while it inhibited the transendothelial migration of the MDA-MB-231 cells (Figures 2G,H). The supernatant from MDA-MB-231 with IL-8 knock down led to decreased endothelial RhoA activation (Suppl. Figures 2J,K), supporting the paracrine role of tumor-derived IL-8 on the endothelium. These results demonstrate that tumor cell-derived IL-8 plays a paracrine regulatory role on transendothelial tumor cell migration.

IL-8 is known to bind and signal through the G-protein coupled receptors CXCR1 and CXCR2 ^47,48^ and RhoA is the downstream target of their activation ^48–51^. SX-682 is a small molecule dual inhibitor of CXCR1 and CXCR2 ^52,53^. Blockade of these receptors with SX-682 blocked IL-8-induced RhoA activation on the endothelial cells (Suppl. Figures 2L,M) and decreased the transendothelial migration of MDA-MB-231 to levels similar to the ones with IL-8 deficiency (Figures 2I,J). Small GTPases, such as RhoA, are activated by guanine nucleotide exchange factors (GEFs), which regulate the cycling from an inactive GDP-bound form to an active GTP-bound form ^35,54^. To identify the GEFs involved in IL-8-mediated RhoA activation we pulled down active GEFs using GST-RhoA^G17A^ as a bait ^32^ and analyzed the activation of four GEFs correlated with RhoA-mediated endothelial permeability: p115-RhoGEF, PDZ-Rho-GEF, LARG and GEF-H1 ^35,55–59^. We found a significant increase in the activation of p115-RhoGEF following 5 min of IL-8 treatment (Figures 2K,L), but not for the other GEFs tested (Figure 2K and Suppl. Figure 2N). To validate the role of p115-RhoGEF in tumor cell transendothelial migration, we knocked down p115-RhoGEF in endothelial cells (Suppl. Figure 2O,P) and performed transmigration experiments with MDA-MB-231 cells. No significant decrease was observed upon endothelial p115-RhoGEF knockdown (Suppl. Figure 2Q,R), suggesting the potential presence of alternative tumor cell-secreted RhoA activators, the participation of other GEFs or insufficient p115-RhoGEF knockdown efficiency. To test the second hypothesis, we treated the endothelial cells with Y16, a GEF inhibitor for LARG, p115-RhoGEF, and PDZ-RhoGEF ^60^. Pretreatment of endothelial cells with Y16 decreased the transendothelial migration of MDA-MB-231 tumor cells (Figure 2M,N). Endothelial GEF inhibition by Y16 was sufficient to block IL-8-induced p115RhoGEF activation (Suppl. Figures 3A,B) and MDA-MB-231 transendothelial cell migration (Suppl. Figures 3C,D). The above demonstrate that tumor cell-derived IL-8 acts in a paracrine manner on endothelial cells, activating RhoA via p115-RhoGEF, inducing endothelial permeability and promoting breast tumor transendothelial migration (Figure 2O).

### High IL-8 expression levels correlate with poor prognosis in human breast tumors

The impact of IL-8 in transendothelial migration prompted us to explore whether IL-8 expression is associated with metastasis-free survival of breast tumor patients. Serum IL-8 levels are considered an independent prognostic marker for overall survival ^61^, while IL-8 mutations have been linked with decreased overall and disease-free survival ^62^. Mining of expression profiling datasets ^38^ revealed that high expression levels of both 77 amino acid (Figure 2P) or 72 amino acid (Figure 2Q) IL-8 correlate with shorter periods of distant metastasis-free survival, even in the subgroup with lymph node-negative breast tumor patients at the time of diagnosis (Suppl. Figures 3E,F). Using the same dataset, similar correlations were not identified for the levels of sTNF-R1 (HR=0.8) or BAFF (HR=0.88) with metastasis-free survival periods of these patients (Suppl. Figures 3G and 3H respectively).

**Figure 3.**
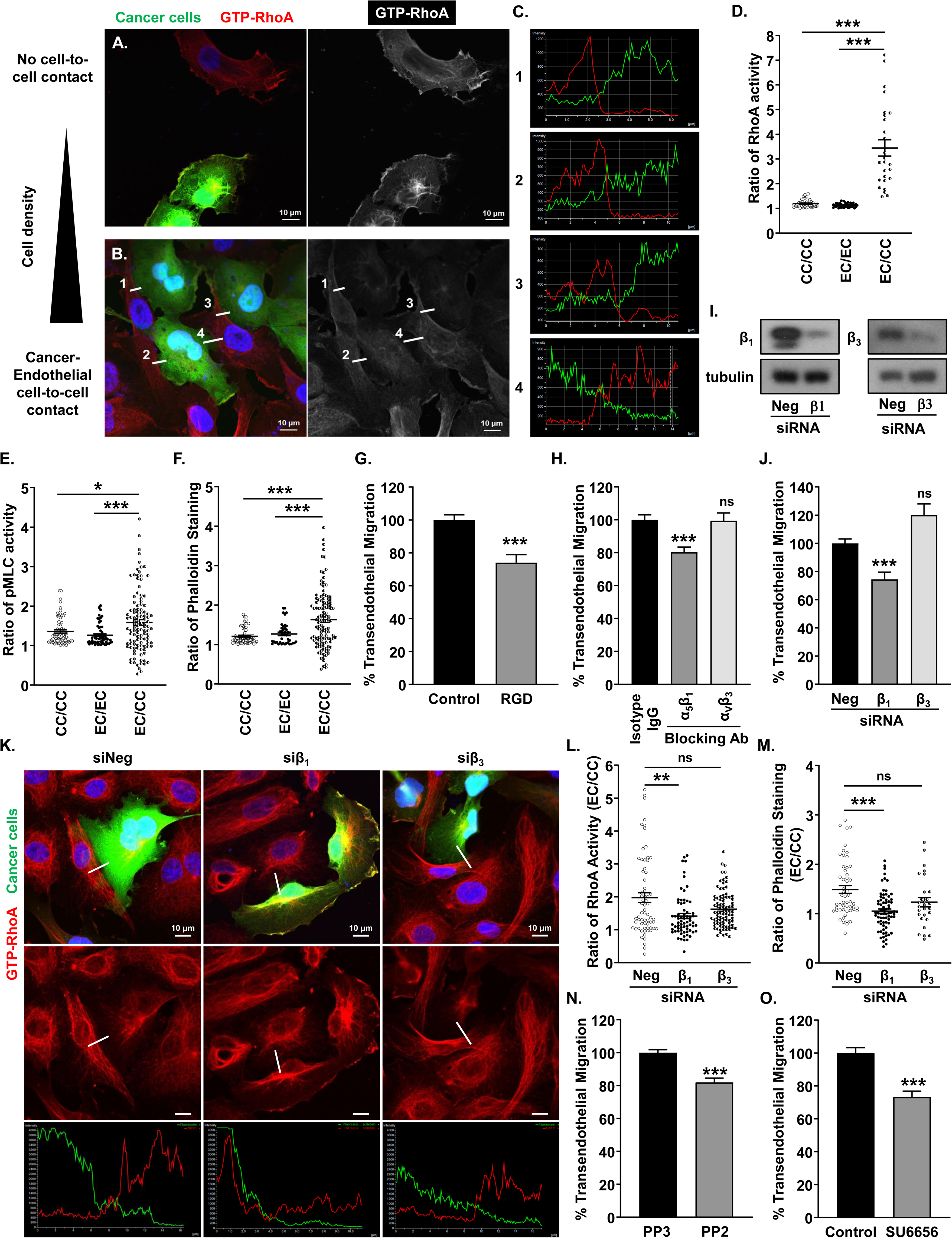
Endothelial RhoA activation during cell-to-cell contact with tumor cells occurs at the focal points and is mediated by endothelial α_5_β_1_ integrin. (A-D) Representative images (A and B) and quantification (C and D) of RhoA activation at the cell-to-cell contact points during co-culture of HUVEC and GFP+ MDA-MB-231 cancer cells; n = 3. (E) Quantification of MLC phosphorylation at the cell-to-cell contact points during co-culture of HUVEC and GFP+ MDA-MB-231 cancer cells; n = 3. (F) Quantification of phalloidin fluorescence intensity at the cell-to-cell contact points during co-culture of HUVEC and GFP+ MDA-MB-231 cancer cells; n = 3. (G) Quantification of transendothelial migration of GFP+ MDA-MB-231 cells after endothelial pretreatment with RGD peptide; n = 3. (H) Quantification of transendothelial migration of GFP+ MDA-MB-231 cells after endothelial pretreatment with Isotype IgG or blocking antibodies against integrins α_5_β_1_ (0.1 µg/ml) and α_V_β_3_ (0.1 µg/ml); n = 3. (I) Representative western blot images of integrin β_1_ and β_3_ knockdown in EAhy926 cells by siRNA treatment (50 nM) at 48 h post-transfection. (J) Quantification of transendothelial migration of GFP+ MDA-MB-231 cells after treatment of the endothelial cells with scramble siRNA (Neg) or siRNA for integrins β_1_ or β_3_; n = 4. (K-L) Representative images (K) and quantification (K and L) of RhoA activation at the cell-to-cell contact points during co-culture of GFP+ MDA-MB-231 cancer cells and HUVEC treated with scramble siRNA (Neg) or siRNA for integrins β_1_ or β_3_; n = 3. (Μ) Quantification of phalloidin fluorescence intensity at the cell-to-cell contact points during co-culture of GFP+ MDA-MB-231 cancer cells and HUVECs treated with scramble siRNA (Neg) or siRNA for integrins β_1_ or β_3_; n = 3. (N) Quantification of transendothelial migration of GFP+ MDA-MB-231 cells after treatment of the endothelial cells with the Src inhibitor (PP2) and the corresponding control (PP3); n = 3. (O) Quantification of transendothelial migration of GFP+ MDA-MB-231 cells after treatment of the endothelial cells with vehicle (Control) or the Src inhibitor (SU6656); n = 4. Data in (D), (E), (F), (G), (H), (J), (L), (M), (N) and (O) represent mean ± SEM. Data in (D), (E), (F), (H), (J), (L) and (M) were analyzed by one-way ANOVA and data in (G), (N) and (O) were analyzed by Student’s unpaired t-test. ns = non significant; *P < 0.05; **P < 0.01; ***P < 0.001.

### Endothelial RhoA activation during cell-to-cell contact with cancer cells occurs at the focal points and is mediated by endothelial α_5_β_1_ integrin

During hematogenous dissemination of circulating tumor cells, the tumor cells come in contact with the endothelial cells in the capillaries or bigger vessels, either through physical entrapment or cell adhesion ^6,63,64^. Physical and biochemical mechanisms have been described during endothelial-cancer cell-to-cell contact ^65–67^ and RhoA activation has been correlated with several mechanisms ^68–70^. After the fast initial arrest of the tumor cell at the endothelium, extravasation is typically completed in one to three days^6^, thus we explored whether tumor-endothelial cell-to-cell contact induces endothelial RhoA activation at the focal points, using a GTP-RhoA-specific antibody ^71^. After overnight co-culture of MDA-MB-231 tumor cells with HUVECs, no distinct difference in RhoA activation was identified in cells without contact (Figure 3A), however, upon direct tumor-endothelial cell-to-cell contact RhoA activation levels were higher in the endothelial (GFP-) than the tumor (GFP+) cell side at the focal points (Figures 3B,C and Suppl. Figure 4A). The ratio of RhoA activation at the focal points of the heterotypic (endothelial/tumor) cell-cell interactions was increased compared to the homotypic (tumor/tumor; endothelial/endothelial) cell-cell interactions (Figure 3D and Suppl. Figure 4A). Induced endothelial RhoA activation seems to be tumor-specific, as it did not occur when the non-tumoral epithelial cell line BEAS-2B was used instead of tumor cells in same experimental setup (Suppl. Figure 4B,C). In agreement with the ratio of RhoA activation levels, the ratios of myosin light chain (MLC) phosphorylation and phalloidin staining were also increased (Figure 3E,F and Suppl. Figure 4D), denoting higher activation of the RhoA downstream signaling within the interacting endothelial cells.

Although tumor integrin expression and activation are mostly known to regulate tumor cell extravasation ^66^, we explored whether endothelial integrin expression and activation play regulatory role in the tumor cell extravasation process. Indeed, treatment of the endothelial monolayer with an RGD (Arg-Gly-Asp) peptide inhibited the transendothelial migration of the MDA-MB-231 cells (Figure 3G and Suppl. Figure 4Ε), denoting integrin involvement. The fibronectin receptors, α_5_β_1_ and α_V_β_3_ integrins regulate important endothelial functions ^72,73^. To explore a potential role of the endothelial α_5_β_1_ and α_V_β_3_ integrins in breast tumor cell transendothelial migration, we used specific blocking antibodies. Endothelial α_5_β_1_, but not α_V_β_3_ blockade inhibited the transendothelial migration of the MDA-MB-231 breast tumor cells (Figure 3H and Suppl. Figure 4F), while co-culture experiments revealed higher β_1_ integrin activation levels in the endothelial than the interacting tumor cells (Suppl. Figure 4G). To explore whether the endothelial β_1_ subunit can regulate tumor cell transendothelial migration, we knocked down the β_1_ and β_3_ subunits in the endothelial cells. Knockdown of the β_1_ subunit inhibited the transmigration of the MDA-MB-231 cells, whereas the knockdown of the β_3_ subunit did not have a negative impact (Figures 3I,J and Suppl. Figures 4H-J). Moreover, endothelial β_1_ knockdown decreased the ratio of heterotypic (endothelial/tumor) RhoA activation at the endothelial-tumor cell focal points (Figures 3K,L), followed by a similar reduction in phalloidin intensity (Figure 3M and Suppl. Figure 4K). To identify whether the downstream integrin signaling affects the transendothelial migration of breast tumor cells, we preincubated the endothelial cells with the Src inhibitors PP2 and SU6656. Endothelial Src inhibition inhibited tumor cell transendothelial migration (Figures 3N,O and Suppl. Figures 4L,M), indicating an important role of integrin outside-in signaling in this process. The tumor cell contact-driven endothelial RhoA activation seems to be an independent mechanism from the paracrine IL-8 effect, as incubation of RGD peptide or blocking antibody against α_5_β_1_ integrin inhibited baseline endothelial RhoA activation, but did not affect IL-8-driven RhoA activation (Suppl. Figure 5A). Given that the paracrine IL-8-driven mechanism and the α_5_β_1_ integrin-mediated one are independent, we wanted to evaluate their combined impact on the metastatic potential using an experimental metastasis *in vivo* model and the murine B16-F10 melanoma cell line, which provides visible lung metastatic lesions. Thus, SX-682 was used to block the IL-8-driven mechanism and the α_5_β_1_ blocking antibody to block the cell-to-cell contact mechanism. Blockade of both pathways via combined treatment led to significantly inhibited metastatic dissemintation (Suppl. Figure 5B-D).

### Endothelial RhoA inhibition decreases cancer cell transendothelial migration

The common signaling mediator of both paracrine and cell-to-cell contact pathways in the endothelial cells is RhoA. To study the functional significance of endothelial RhoA on cancer cell transendothelial migration, we pharmacologically blocked endothelial RhoA by pretreatment of the endothelial monolayer with the Rho GTPase inhibitor C3 toxin ^17,29^. Endothelial RhoA inhibition significantly reduced the transmigration of MDA-MB-231 cancer cells across the endothelial monolayer of both EAhy926 (Figure 4A) and HUVECs (Figure 4B). Contrary to the genetic instability of the tumor cells, the tumor microenvironment, and in this case the endothelial cells, should provide a more predictable target for pharmaceutical intervention, independent of the driving mutations of the cancer type ^3,4^ and potentially applicable to different solid tumors. For this, and in order to assess the impact of endothelial RhoA inhibition on tumor metastasis in an immune-competent *in vivo* microenvironment, we selected three murine syngeneic models, EO771 breast tumor, B16-F10 melanoma, and Lewis Lung Carcinoma (LLC) cell lines. We first explored whether the secretome from these cell lines can activate RhoA in the endothelial cells. Starvation supernatant from all tumor cell lines activated endothelial RhoA (Suppl. Figures 6A-C). Endothelial RhoA blockade with C3 toxin significantly inhibited the transendothelial migration of murine syngeneic EO771 breast cancer cells (Figure 4C), as well as B16-F10 (Figure 4D) and LLCs (Figure 4E), demonstrating a uniform role of endothelial RhoA on transendothelial tumor cell migration of solid tumors. Moreover, C3-toxin pretreatment was sufficient to block TEM of MDA-MB-231 even when stimulated with higher than normally secreted amount of IL-8 (50 ng/ml) (Suppl. Figure 6D,E). Interestingly, the transendothelial migration of non-cancerous, but migratory NIH-3T3 cells was mostly unaffected by endothelial RhoA inhibition by C3 toxin (Suppl. Figure 6F,G)

**Figure 4.**
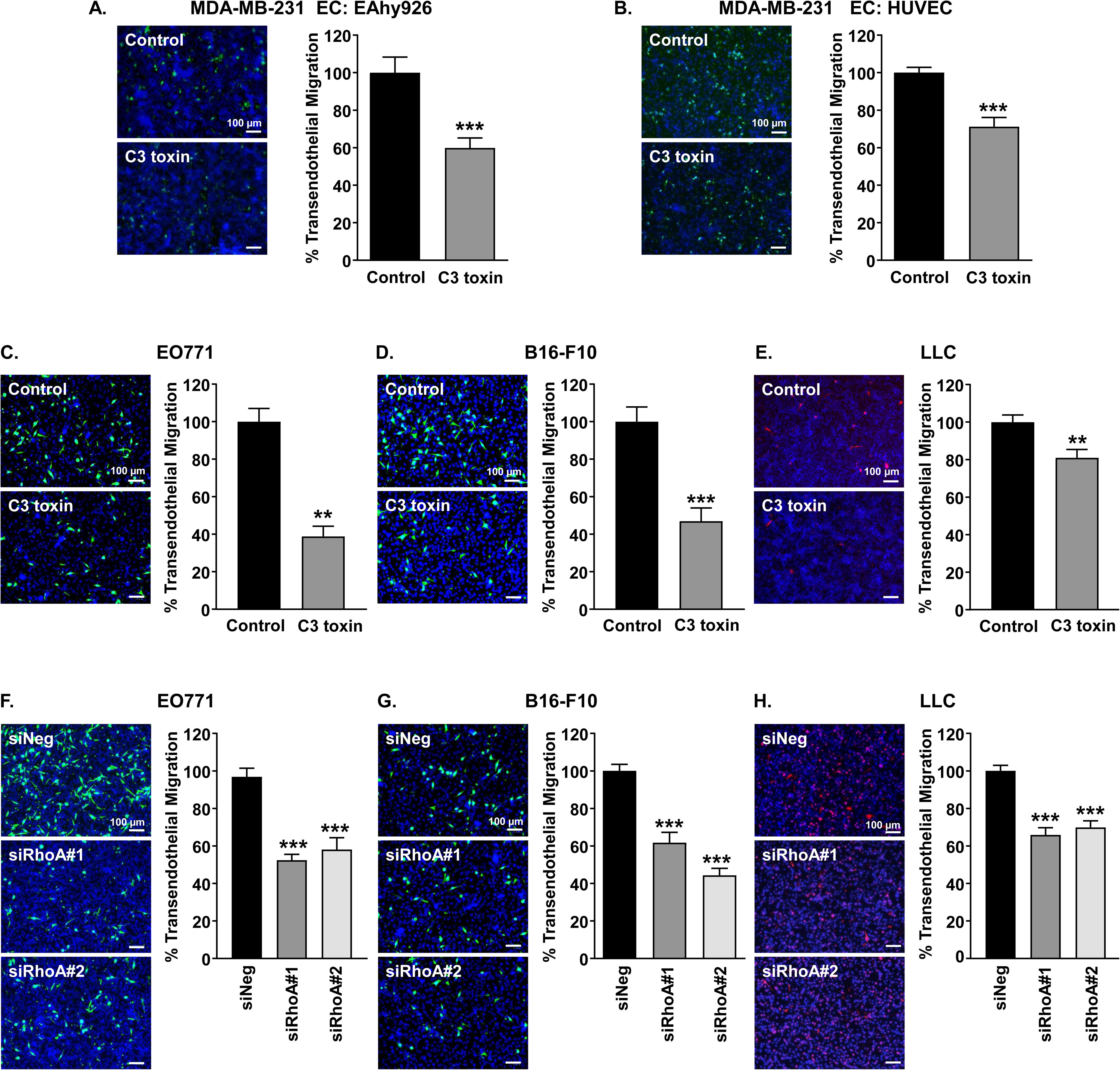
Endothelial RhoA inhibition blocks tumor cell transendothelial migration. (A and B) Representative images and quantification of transendothelial migration of GFP+ MDA-MB-231 cells through an EAhy926 (A) and HUVEC (B) endothelial monolayer pretreated with vehicle or C3 toxin (20 ng/ml) (nuclei = blue); A, n = 3; B, n= 4; Scale bars, 100 µm. (C-E) Representative images and quantification of transendothelial migration of RFP+ or green fluorescent dye-labeled murine EO771 (C), B16-F10 (D) and LLC (E) cancer cells, through the EAhy926 endothelial monolayer pretreated with vehicle or C3 toxin (20 ng/ml) (nuclei = blue); C, n = 2; D, n = 4; E, n = 3; Scale bars, 100 μm. (F-H) Representative images and quantification of trans-endothelial migration of RFP+ or green fluorescent dye-labeled murine EO771 (F), B16-F10 (G) and LLC (H) tumor cells through siNeg- or siRhoA-transfected (50 nM) EAhy926 endothelial monolayer (nuclei = blue); F, n = 2; G, n = 3; H, n = 4; Scale bars, 100 μm. Data in (A), (B), (C), (D), (E), (F), (G) and (H) represent mean ± SEM. Data in (A), (B), (C), (D) and (E) was analyzed by Student’s unpaired t-test and data in (F), (G) and (H) was analyzed by one-way ANOVA. **P < 0.01; ***P < 0.001.

Although C3 toxin is a widely used pharmacological inhibitor of RhoA, it is known to block activation of other Rho isoforms, including RhoA, RhoB, RhoC and RhoE ^74^. Therefore, we selectively knocked down endothelial RhoA with specific siRNAs (Suppl. Figures 6H,I). For both siRNA sequences used, knock-down of RhoA in the endothelial cells resulted in a decrease of cancer cell migration for MDA-MB-231 (Suppl. Figures 6J,K), EO771 (Figure 4F), B16-F10 (Figure 4G) and LLC (Figure 4H) tumor cells.

### Endothelial RhoA deficiency prevents lung metastasis of syngeneic murine cancer cells

To pinpoint the role of endothelial RhoA in metastatic outcome *in vivo*, we used tamoxifen-inducible endothelial-specific RhoA deficient mice ^17^. Endothelial RhoA deficiency is induced by the tamoxifen-inducible *Cdh5-CreERT*^2^ promoter (Figure 5A). Excision efficiency was monitored with the tdTomato-green fluorescent protein (GFP) reporter ^75^, where GFP-expressing (green) endothelial cells can be visualized upon CRE-mediated gene recombination (Figure 5B). Western blot analysis of isolated lung ECs from tamoxifen-treated RhoA^iΔΕC^ mice showed no RhoA expression, and GFP expression instead (Figure 5C). The lung is a frequent site of metastases from extrathoracic tumors and the second most common site of metastasis for breast cancer ^76–78^, therefore we assessed the number of lung micrometastases as a quantitative marker for the evaluation of the metastatic potential of the *in vivo* models.

**Figure 5.**
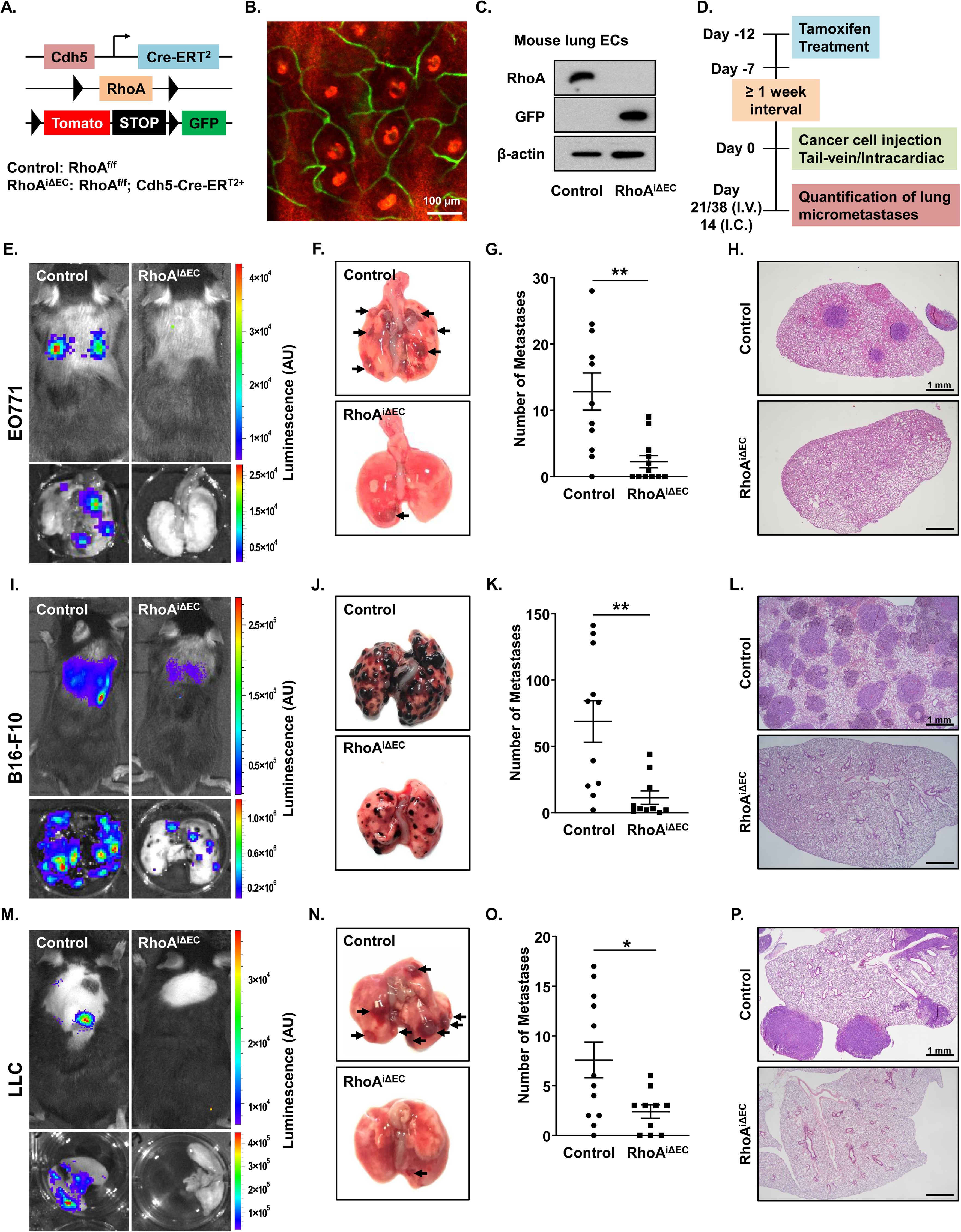
Endothelial RhoA deficiency prevents lung metastases of syngeneic murine tumor cells. (A) Schematic representation of endothelial-specific conditional RhoA-deficient (RhoA^iΔEC^) mice with Tomato-GFP reporter. (B) Representative confocal image of mouse ear of Tamoxifen-treated RhoA^f/f^; Tomato-GFP^f/f^; *Cdh5-CreERT^2^+* mouse showing excision efficiency driven by the Cdh5-CreERT^2^ promoter (GFP+ capillaries surrounding the hair follicles/intense red spots). Scale bars, 100 μm. (C) Lung endothelial cells, isolated from RhoA^f/f^; Tomato-GFP^f/f^ mice without (Control) or with *Cdh5-CreERT*^2^ promoter (RhoA^iΔEC^) were treated with Tamoxifen for five consecutive days and analyzed for RhoA, GFP and actin expression with western blot. (D) Schematic diagram of experimental metastasis experiments. Mice were given 2 mg Tamoxifen/mouse/day intraperitoneally (IP) for five consecutive days. At least one week after the last Tamoxifen treatment, 1×10^5^ cancer cells were administered either through intravenous (I.V.) or intra-cardiac (I.C.) route. At the indicated days, the experiments were terminated and the number of metastases was evaluated from the dissected lungs. (E-H) Evaluation of metastatic outcome of EO771 syngeneic breast cancer cell line in RhoA^iΔEC^ mice and littermate controls (IV administration). (E) Representative IVIS images of the mice and dissected lungs. (F) Representative macroscopic appearance of the dissected lungs (arrows denote metastases). (G) Quantification of number of metastases per mouse; Control, n = 11; RhoAiΔEC, n = 12. (H) Representative H&E lung sections. Scale bars, 1 mm. (I-L) Evaluation of metastatic outcome of B16-F10 syngeneic melanoma cell line in RhoA^iΔEC^ mice and littermate controls (IV administration). (I) Representative IVIS images of the mice and dissected lungs. (J) Representative macroscopic appearance of the dissected lungs. (K) Quantification of number of metastases per mouse; Control, n = 11; RhoA^iΔEC^, n = 10. (L) Representative H&E lung sections. Scale bars, 1 mm. (M-P) Evaluation of metastatic outcome of LLC syngeneic lung cancer cell line in RhoA^iΔEC^ mice and littermate controls (IV administration). (M) Representative IVIS images of the mice and dissected lungs. (N) Representative macroscopic appearance of the dissected lungs (arrows denote metastases). (O) Quantification of number of metastases per mouse; Control, n = 12; RhoA^iΔEC^, n = 10. (P) Representative H&E lung sections. Scale bars, 1 mm. Data in (G), (K) and (O) represent mean ± SEM and were analyzed by Student’s unpaired t-test. *P < 0.05; **P < 0.01.

To test the role of endothelial RhoA in metastasis in an immunocompetent environment we used syngeneic triple-negative breast cancer cells EO771, which have well-characterized metastatic potential ^79,80^, and tumor cells were administered in the caudal vein (intravenous administration or I.V. model) (Figure 5D). Endothelial RhoA deficiency decreased the metastatic potential of intravenously administered EO771 breast tumor cells (Figures 5E-H). The mice with endothelial RhoA deficiency (RhoA^iΔEC^) presented decreased luciferase signal in the lung area than the littermate controls, demonstrated by reduced bioluminescence (Figure 5E), fewer visible lung metastases (Figures 5F,G) and fewer tumoral areas identified by H&E staining (Figure 5H).

One of the advantages of targeting the tumor microenvironment for tumor growth and metastasis inhibition is the elimination of the differential response and resistance due to heterogeneity within the same tumor, which determines the therapeutic outcome ^81^, as well as applicability in diverse tumor types ^82^. To identify whether endothelial RhoA deficiency has potential for a universal anti-metastatic treatment, we performed experimental metastasis experiments, assessing mostly the extravasation part of the metastatic process, with diverse tumor types, and used two other syngeneic tumor lines for the C57BL/6J background, B16-F10 melanoma and LLC carcinoma. Both cell lines are frequently used for experimental metastasis experiments and are known to colonize in the lungs following systemic inoculation ^83–85^. When injected intravenously, both B16-F10 and LLCs metastasized in the lungs. In line with our results of the murine breast tumor model, endothelial RhoA-deficiency reduced metastatic burden in the lungs for both the B16-F10 (Figures 5I-5L) and the LLC cells (Figures 5M-5P), indicating that endothelial RhoA activation plays significant role during tumor cell transmigration and metastasis, irrespectively of the solid tumor type.

Tumor cells injected via lateral tail vein encounter pulmonary capillary beds first, whereas the introduction of the cells to arterial circulation via the left ventricle of the heart (intracardiac administration) disseminates the cells throughout the body ^86,87^. Thus, to identify whether the route of tumor cell administration changes the impact of RhoA deficiency on metastasis and to validate our *in vivo* findings with another model, we performed experimental metastasis experiments where the tumor cells were injected into the left ventricle of the heart (intracardiac or I.C. model) (Figure 5D). Despite the equal number of tumor cells being injected, the intracardiac model developed metastases earlier than the intravenous one. Nevertheless, endothelial RhoA deficiency reduced the metastatic potential of B16-F10 (Suppl. Figures 7A-D) and LLC (Suppl. Figures 7E-5H) cells. We also assessed the role of endothelial RhoA in a spontaneous metastasis model using EO771 cells. The numbers of lung metastases (Suppl. Figures 7I-J) and circulating tumor cells (Suppl. Figures 7K-L) were significantly reduced, suggesting that endothelial RhoA is crucial also for the intravasation step of metastasis. Taken together, these data demonstrate that the role of endothelial RhoA on cancer cell transendothelial migration and metastasis is significant and independent of the site of tumor dissemination or the tumor type.

### Pharmacological blockade of Rho Kinase (ROCK) inhibits tumor cell transendothelial migration

Upon activation, RhoA activates Rho-associated protein kinase (ROCK) ^88,89^. Previous studies have shown the stabilization of endothelial barrier function through ROCK inhibition in various pathological conditions ^17,18,90–92^. We therefore assessed the effects of endothelial ROCK inhibition on endothelial cells in tumor cell transendothelial migration. Pretreatment of endothelial cells with ROCK inhibitor Y-27632 ^93^ and the clinically relevant inhibitor Fasudil ^94^ reduced the transmigration of MDA-MB-231 cells across monolayers of both EAhy926 cells (Figure 6A) and HUVECs (Figure 6B). Pretreatment with both ROCK inhibitors blocked the transmigration of all murine tumor cell lines examined: EO771 (Figure 6C), B16-F10 (Fig 6D) and LLC (Figure 6E).

**Figure 6.**
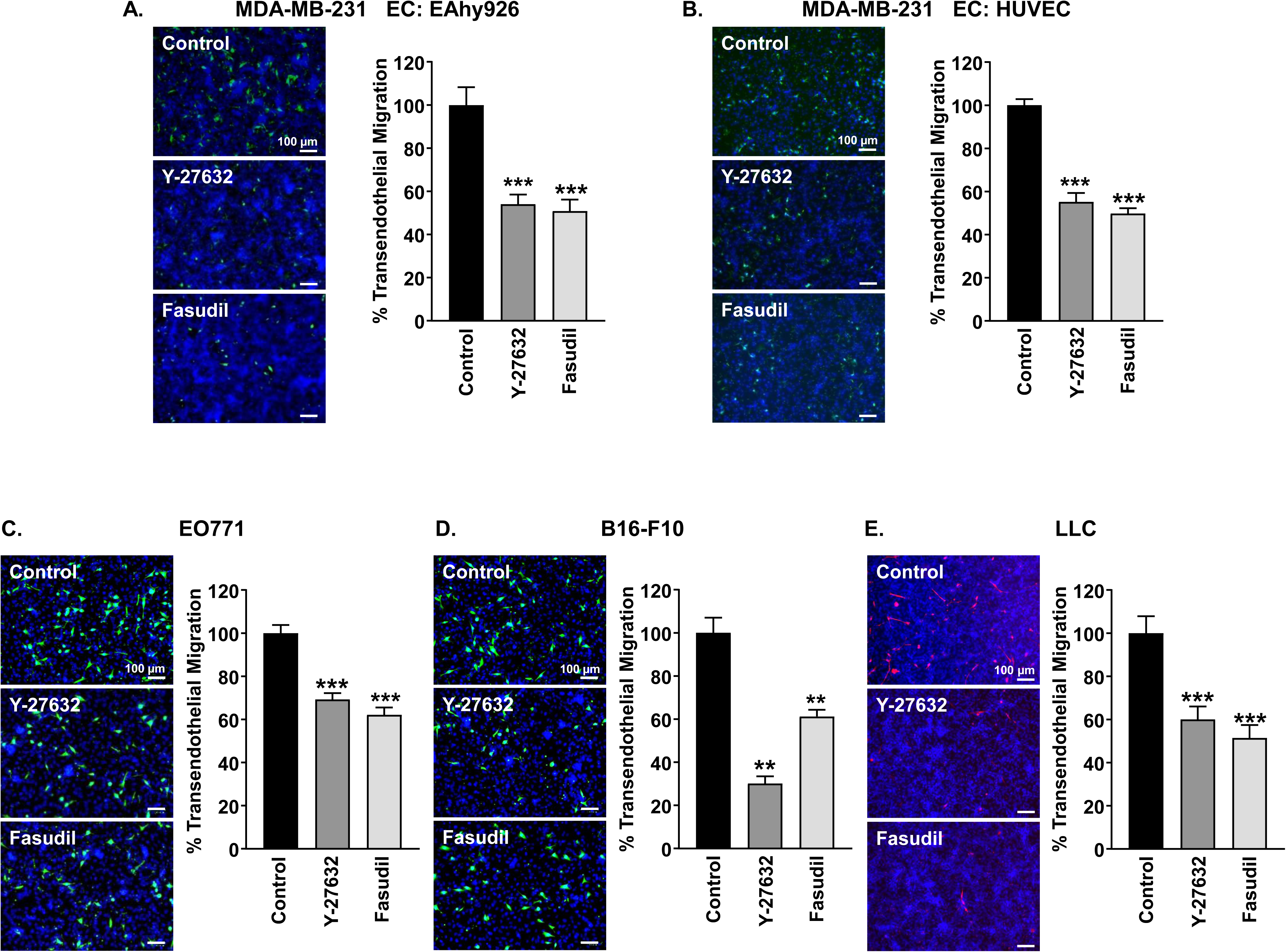
Pharmacological blockade of Rho kinase (ROCK) inhibits tumor cell trans-endothelial migration. (A and B) Representative images and quantification of transendothelial migration of GFP+ MDA-MB-231 cells through an EAhy926 (A) and HUVEC (B) endothelial monolayer pretreated with vehicle, Y-27632 (10 μM) or Fasudil (10 μΜ) (nuclei = blue); A: n = 3 B: n= 3. Scale bars, 100 µm. (C-E) Representative images and quantification of transendothelial migration of green fluorescent dye-labelled or RFP+ murine EO771 (C), B16-F10 (D) and LLC (E) cancer cells, through an EAhy926 endothelial monolayer pretreated with vehicle, Y-27632 (10 μM) or Fasudil (10 μΜ) (nuclei = blue); C, n = 2; D, n = 3; E, n = 3. Scale bars, 100 μm. Data in (A), (B), (C), (D) and (E) represent mean ± SEM and was analyzed by one-way ANOVA. **P < 0.01; ***P < 0.001.

### Treatment with clinically relevant ROCK inhibitor blocks metastatic potential of murine and human tumor cells

The role of ROCK in tumor transformation, growth, invasion and angiogenesis has been well documented and the ROCK inhibitors Y27632 and Fasudil are widely used ^95^. Here, with a focus on the endothelial RhoA-ROCK pathway, we explored the effects of Fasudil in the metastatic fate of the syngeneic murine cancer cells EO771, B16-F10 and LLC. Since the circulating tumor cells do not spend much time in the circulation and the initial entrapment in the capillaries, followed by transmigration, plays a crucial role in the metastatic outcome ^6,64^, we initiated Fasudil treatment one day before tumor cell inoculation. Daily intraperitoneal administration of Fasudil (20 mg/kg) significantly reduced the metastatic load in the lungs for breast tumor (EO771; Figures 7A-D), melanoma (B16-F10; Figures 7E-H) and lung carcinoma cells (LLC; Figures 7I-L). Similar results were also obtained through intracardiac administration of the tumor cells, tested for B16-F10 (Suppl. Figures 8A-D) and LLC (Suppl. Figures 8E-H) metastasis.

**Figure 7.**
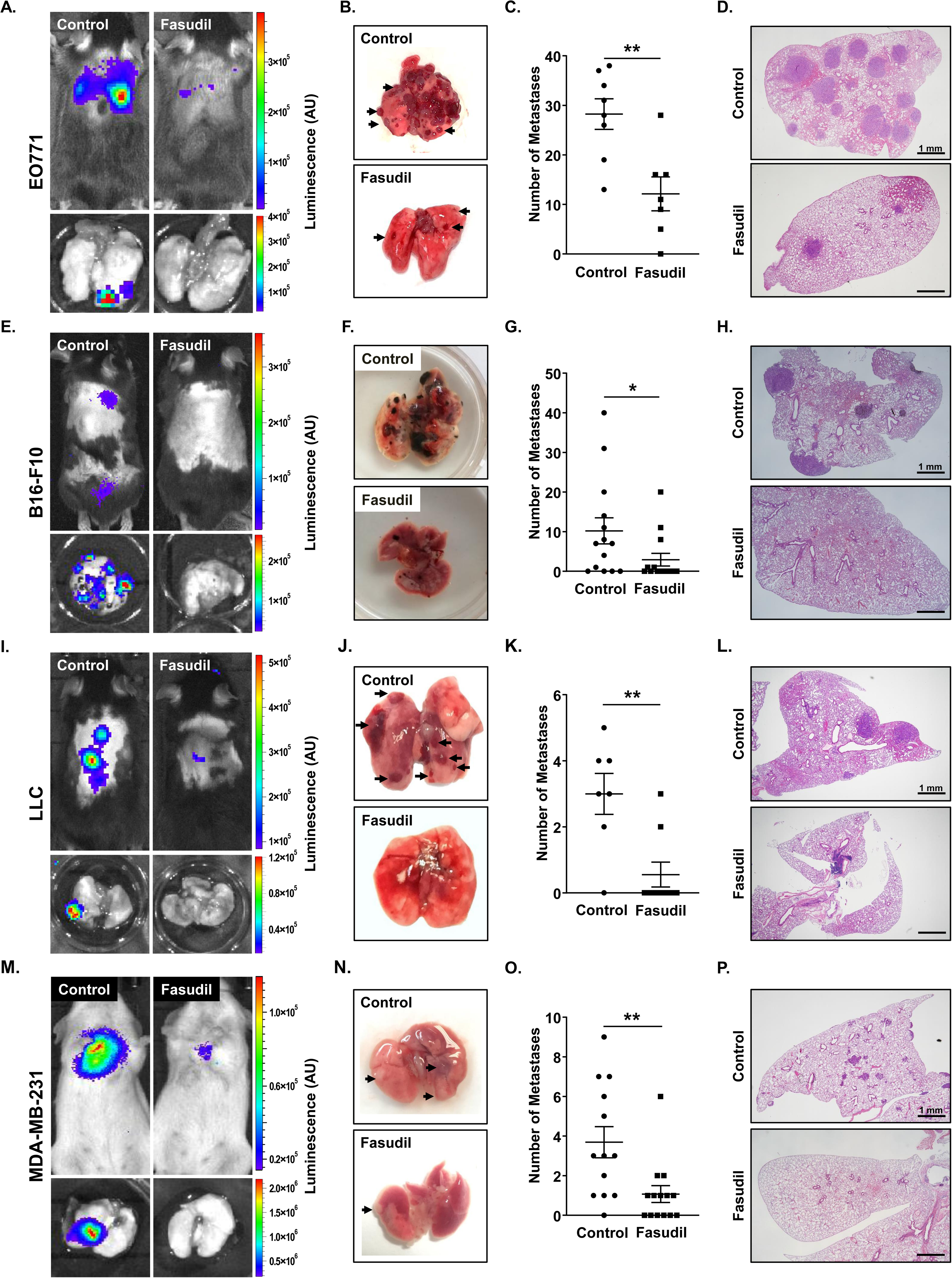
Treatment with clinically relevant ROCK inhibitor blocks metastatic potential of murine and human tumor cells. (A-D) Evaluation of metastatic outcome of EO771 syngeneic breast cancer cell line (I.V.) in mice treated daily with vehicle or Fasudil (20 mg/kg, I.P.). (A) Representative IVIS images of the mice and dissected lungs. (B) Representative macroscopic appearance of the dissected lungs (arrows denote metastases). (C) Quantification of number of metastases per mouse; Control, n = 8; Fasudil, n = 7. (D) Representative H&E lung sections. Scale bars, 1 mm. (E-H) Evaluation of metastatic outcome of B16-F10 syngeneic melanoma cell line (I.V.) in mice treated daily with vehicle or Fasudil (20 mg/kg; I.P.). (E) Representative IVIS images of the mice and dissected lungs. (F) Representative macroscopic appearance of the dissected lungs. (G) Quantification of number of metastases per mouse; Control, n = 14; Fasudil, n = 14. (H) Representative H&E lung sections. Scale bars, 1 mm. (I-L) Evaluation of metastatic outcome of LLC syngeneic lung cancer cell line (I.V.) in mice treated daily with vehicle or Fasudil (20 mg/kg; I.P.). (I) Representative IVIS images of the mice and dissected lungs. (J) Representative macroscopic appearance of the dissected lungs (arrows denote metastases). (K) Quantification of number of metastases per mouse; Control, n = 7; Fasudil, n = 9. (L) Representative H&E lung sections. Scale bars, 1 mm. (M-P) Evaluation of metastatic outcome of MDA-MB-231 human breast cancer cell line (I.V.) in immune compromised mice treated daily with vehicle or Fasudil (20 mg/kg; I.P.). (M) Representative IVIS images of the mice and dissected lungs. (N) Representative macroscopic appearance of the dissected lungs (arrows denote metastases). (O) Quantification of number of metastases per mouse; Control, n = 13; Fasudil, n = 14. (P) Representative H&E lung sections. Scale bars, 1 mm. Data in (C), (G), (K) and (O) represent mean ± SEM and were analyzed by Student’s unpaired t-test. *P < 0.05; **P < 0.01.

We further assessed the effect of Fasudil priming on the metastatic outcome. Fasudil administration before inoculation of tumor cells showed better outcome than treating the mice after (Suppl. Figures 8I-K), demonstrating that Fasudil impacts the extravasation step. To verify Faudil’s impact on extravasation, we performed a colony formation assay, where Fasudil-treated lungs generated significantly fewer antibiotic-resistant colonies, thus extravasated tumor cells than the control ones (Suppl. Figures 8L-M). Although the anti-tumor effect of Fasudil cannot be ignored, these data highlights the significance of ROCK inhibition in tumor cell transendothelial migration.

Several studies have reported positive results of Fasudil using human xenograft (solid tumors and cell lines) models ^95^, while a study reported that Fasudil priming can delay the metastatic onset and enhance the chemotherapeutic outcome in pancreatic tumors ^96^. In triple-negative breast cancer, Fasudil inhibits tumor growth and cell migration ^97,98^ but the effects of Fasudil in the metastasis of triple-negative human breast cancer *in vivo* is unknown. To explore the possibility of repurposing Fasudil as an anti-metastatic drug, we injected the MDA-MB-231 cells into SCID mice upon Fasudil treatment (20 mg/kg). Fasudil treatment blocked the metastatic burden, observed by the decreased number of tumor nodules in the lungs of the Fasudil-treated group (Figures 7M-7P).

The history of Fasudil in the clinic has proven it is well-tolerated in patients without significant adverse effects ^99,100^. In our experimental animals, no differences in mouse weight were observed among the Fasudil- and vehicle-treated mice (Suppl. Figures 9A-H), presenting no toxicity indications.

## Discussion

Given its significance on tumor metastasis and the increased genetic stability compared to the tumor itself ^1,3^, the vasculature is an important and more predictable target for pharmaceutical intervention. Normalization of the vasculature both in the tumoral area and in the metastasis-target organs decreases metastatic incidence and improves immunotherapeutic efficacy ^101^. Our findings demonstrate that endothelial RhoA activation is an important driving event for transendothelial migration and metastatic outcome in solid tumors (Figure 8). Although RhoA has been extensively studied in tumor cells its role remains controversial ^102,103^, whereas the physiology of RhoA in the vasculature is well established ^104,105^, providing a safe target for vascular normalization.

**Figure 8.**
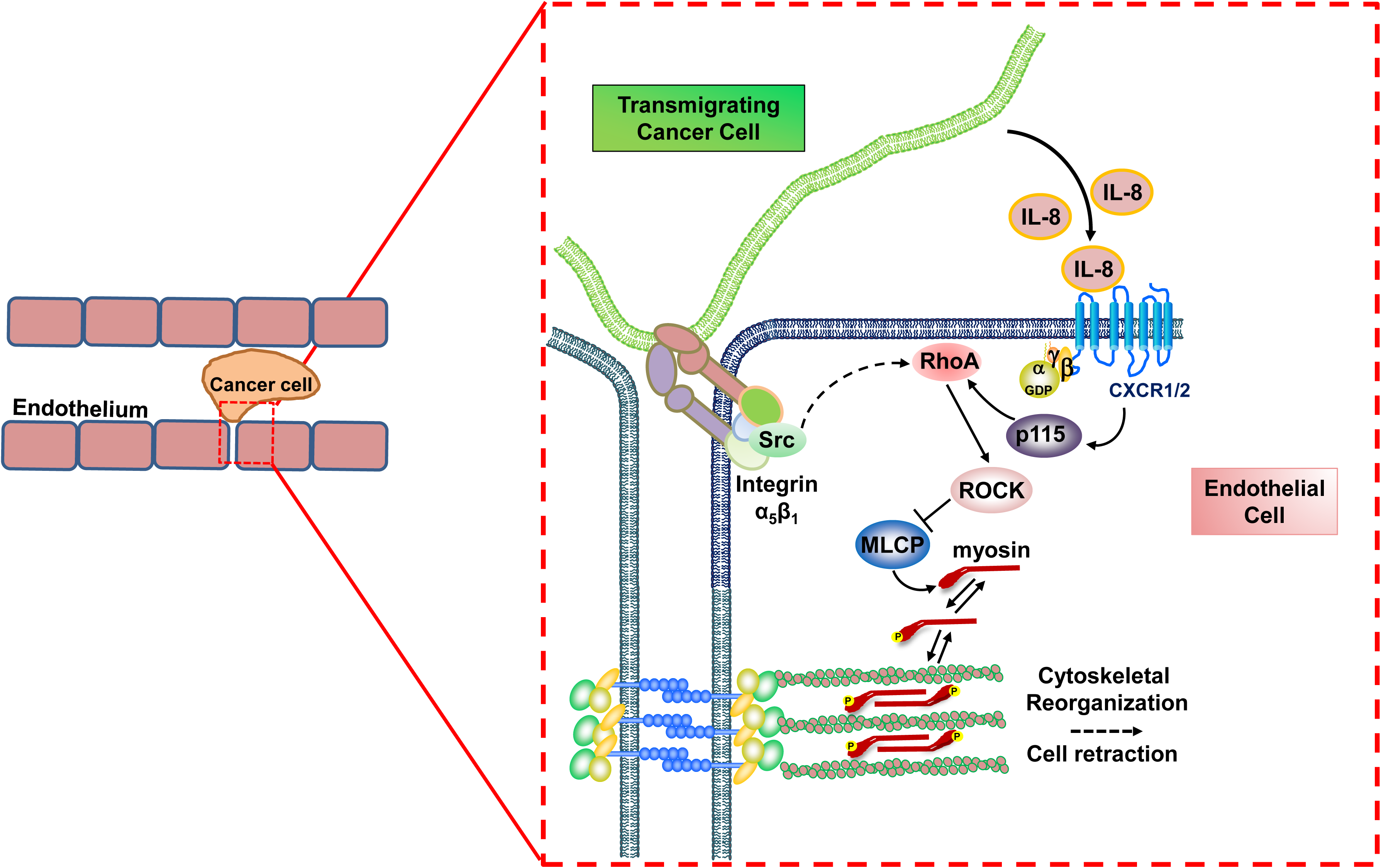
Schematic diagram illustrating the two mechanisms of tumor cell-driven endothelial RhoA activation during transendothelial migration.

GPCR and tyrosine kinase receptor ligands induce endothelial permeability by activating the endothelial RhoA pathway ^15,17^. Primary tumors induce formation of sites of focal hyperpermeability in the lungs prior to cancer cell homing ^80^. IL-8 is a known inducer of endothelial permeability ^51^ and tumor cells with higher metastatic potential have higher IL-8 expression ^106^. Here we report differences in IL-8 secretion levels from breast tumor cell lines with diverse metastatic potential and performed gain- and loss-of-function experiments to evaluate its functional role on breast tumor transmigratory potential. In benign prostatic hyperplasia cells IL-8 has been shown to activate the RhoA/ROCK pathway ^49^. In endothelial cells, IL-8 binds to CXCR1 and CXCR2, leading to cytoskeletal reorganization and upregulates RhoA expression ^48,50^. IL-8-induced endothelial permeability has been also attributed to CXCR1 and CXCR2 binding with VEGFR2, leading to its transactivation in a VEGF-independent manner. The receptor complex formation is mediated by Src, and VEGFR2 inhibition blocks IL-8-induced RhoA activation and permeability ^51^. It is of interest that IL-8 knockdown did not abrogate the transendothelial migration of MDA-MB-231 cells to the level that ROCK inhibition did, which implies the presence of several other mediators, not detected in this study, and the role of alternative cell-to-cell contact mechanisms. However, it provides a proof-of-principle of a potential paracrine mechanism of endothelial RhoA activation during transendothelial cancer cell migration.

Endothelial retraction is a mechanism related with circulating cancer cell transendothelial migration upon their entrapment in capillaries ^1,107^, and we and others have shown that cell retraction is mediated by RhoA activation ^35,108,109^. In the co-culture experiments, we identified increased RhoA activation in the endothelial part of the cell-to-cell contact site, and integrins seem to be involved in RhoA activation and the transmigration process. Integrins are important mediators of cancer metastasis. Integrin function mediates the attachment of circulating tumor cells to the vasculature, their extravasation and dormancy or proliferation to develop metastases^66^. In breast tumor cells, increased β_1_ integrin expression was correlated with increased tumor cell migration ^110^ and tumor cell β_1_ integrin is critical in establishing the adhesion of the circulating tumor cells with the endothelium before and during extravasation ^67,111^. Here we show that the endothelial cells present higher integrin β_1_ activation levels than the surrounding tumor cells and that integrin β_1_ activation correlates with endothelial RhoA activation and regulates the transmigration of the tumor cells. Integrin activation could be due to either receptor or ligand interaction from the tumor cell side or tumor-driven mechanotransduction mechanisms. Force-induced membrane deformation activates integrins and rigid matrix activates β_1_ integrin and induces RhoA-ROCK signaling-mediated contractility in tumor cells and fibroblasts ^112–114^, while the same signaling pathway is activated by Ang2 in endothelial cells ^115,116^. It was recently shown that targeted therapy against α_5_β_1_ integrin reduces liver metastases in a preclinical model ^117^ and clinical samples from breast tumor patients have shown higher α_5_β_1_ integrin expression on tumor-associated endothelial cells ^118^. The fact that α_5_β_1_ integrin levels are correlated with IL-8 expression ^118^ and that cytokine stimulation increases the expression levels of both α_5_ and β_1_ integrin subunits ^119^ could denote converging of paracrine and cell-to-cell contact mechanisms, however our findings denote that does not seem to be the case, at least in terms of endothelial RhoA activation.

It was previously shown that knockdown for endothelial RhoA in HUVECs abrogates prostate tumor cell migration ^120^, which shown here with human and murine breast, lung tumor and melanoma cell lines. Focusing on the tumor-endothelial cell interaction, the *in vitro* part of this study did not investigate a complex but important parameter: the role of other cells in the tumor microenvironment, such as fibroblasts or macrophages, which was later addressed *in vivo* through the incorporation of syngeneic tumor models. Fibroblasts communicate with tumor cells mediating their growth and aggressiveness by growth factor, cytokine and chemokine secretion, while breast tumor cell transendothelial migration is also dependent on tumor cell-macrophage communication ^41,121^.

In neoplastic conditions, the studies on Rho GTPase signaling have been mostly focused on tumor cells. Since RhoA and ROCK mutations have been detected in several tumors, ROCK inhibitors have been preclinically proved to be effective in tumor inhibition and resistance without compromising immune functions ^95,122,123^. Fasudil is clinically used for subarachnoid hemorrhage^124^. Here, we demonstrated that Fasudil-induced inhibition of the endothelial RhoA pathway abrogated tumor cell transendothelial migration *in vitro* and metastasis *in vivo*, highlighting the inhibition of the endothelial RhoA pathway as an efficient, common target of the tumor microenvironment for anti-metastatic therapy.

## KEY RESOURCES TABLE

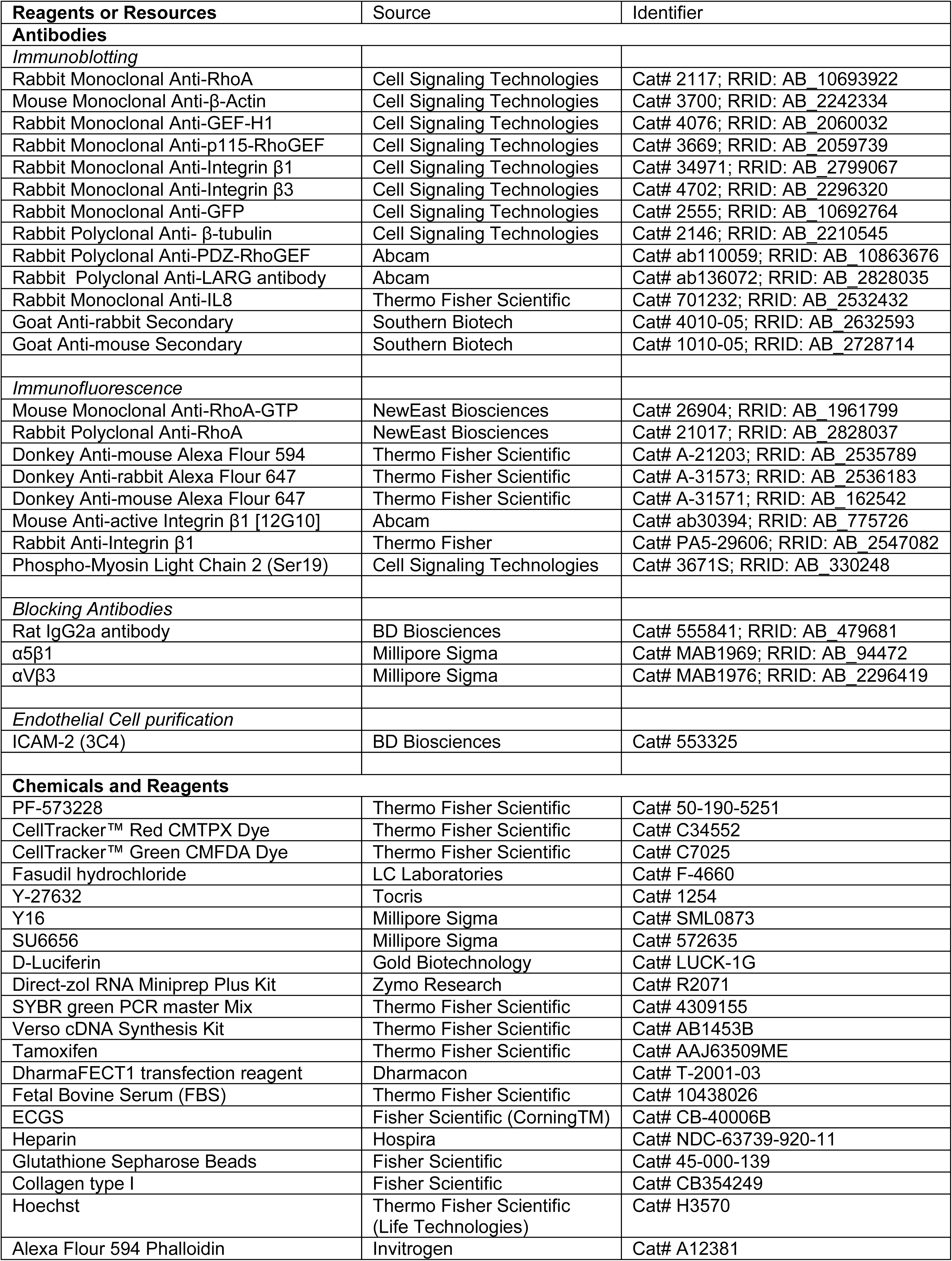
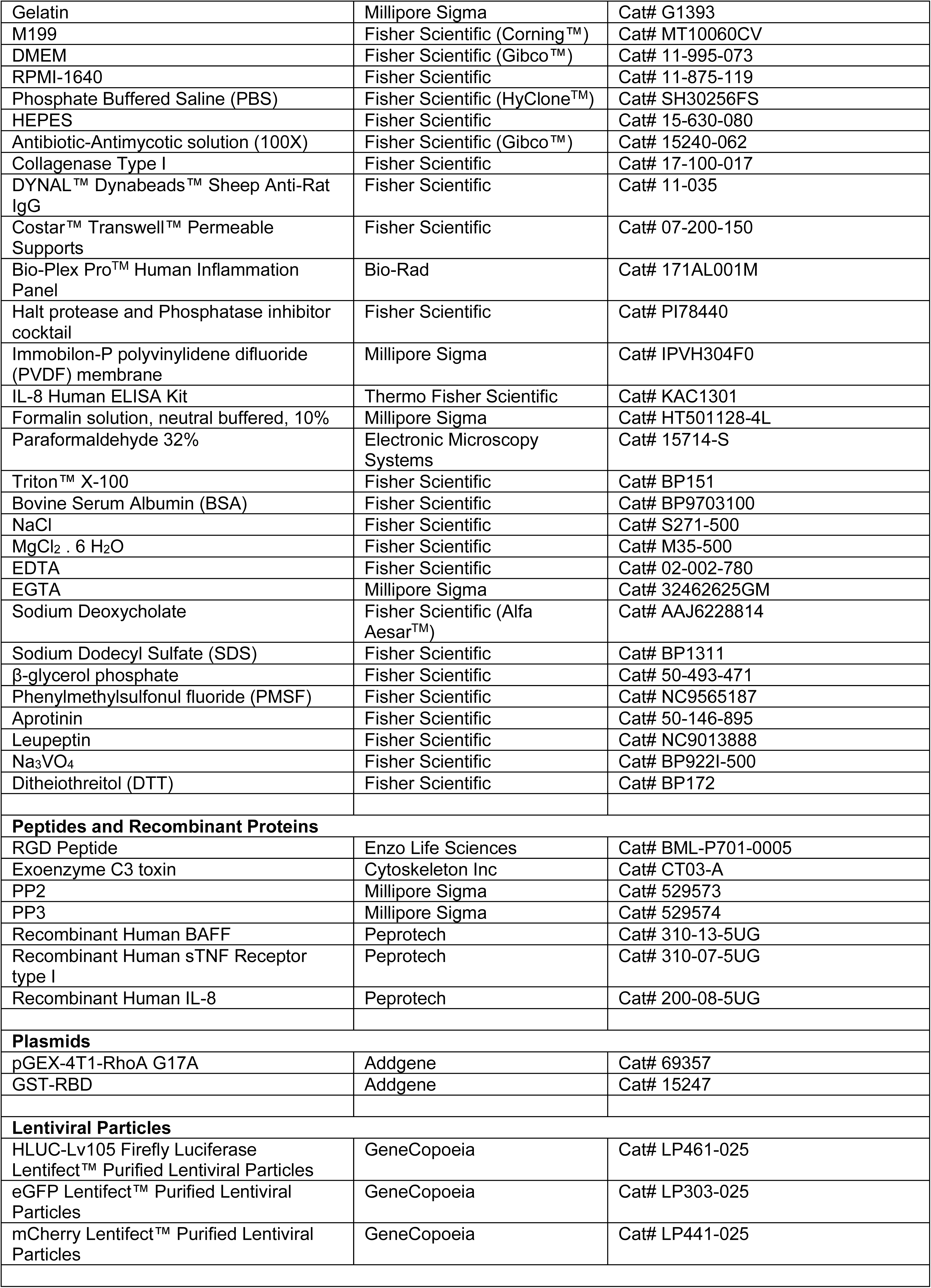
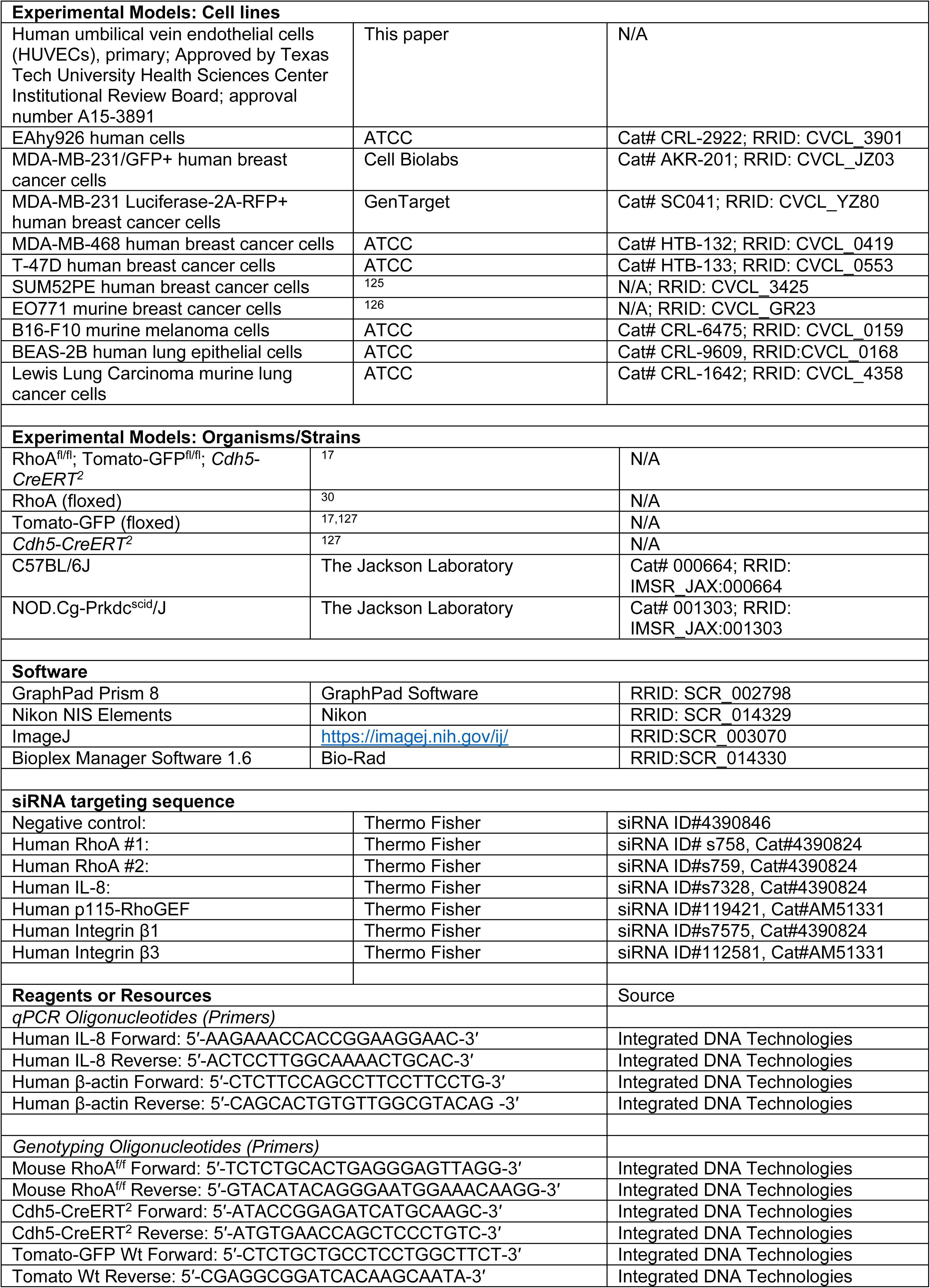
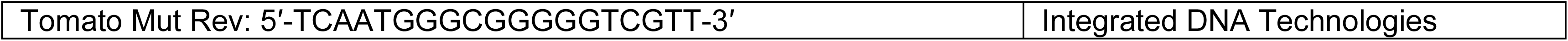

## Acknowledgments

We thank Ralf Adams (MPI Münster, Germany) for providing Chd5-Cre-ERT2 mice and the members of the TTUHSC animal facility in Amarillo for their support. This work was supported in part by the Texas Tech University Health Sciences Center (TTUHSC) School of Pharmacy Office of the Sciences grant, by the Division of Intramural Research of the NIAID, NIH, and by the Hellenic Foundation for Research and Innovation (00376). The funders had no role in study design, decision to write, or preparation of the manuscript.

## Author Contributions

Conceptualization, M.S.S. and C.M.M.; Methodology, M.S.S., J.H.P., M.O. and S.T.; Validation, F.T.Z. and R.G.A.; Investigation, M.S.S., F.T.Z., M.O., R.G.A., C.Z., S.H., J.V.B., S.T. and C.M.M.; Writing – Original Draft, M.S.S. and C.M.M.; Writing – Review & Editing, F.T.Z., C.L.D., S.H., J.V.B., M-H.W., M.M.M., S.K.S., Y.Z., J.S.G., J.H., U.B., S.T.; Resources, P.T., C.L.D., M-H.W., M.M.M., S.K.S., Y.Z., J.S.G., M.S.L., D.Y.M., J.A.Z.; Supervision, J.H., U.B., and C.M.M.; Funding Acquisition: C.M.M.

## References

1 Massague, J. & Obenauf, A. C. Metastatic colonization by circulating tumour cells. Nature 529, 298–306 (2016). 10.1038/nature17038

2 Dillekas, H., Rogers, M. S. & Straume, O. Are 90% of deaths from cancer caused by metastases? Cancer Med 8, 5574–5576 (2019). 10.1002/cam4.2474

3 Quail, D. F. & Joyce, J. A. Microenvironmental regulation of tumor progression and metastasis. Nat Med 19, 1423–1437 (2013). 10.1038/nm.3394

4 Hanahan, D. & Weinberg, R. A. Hallmarks of cancer: the next generation. Cell 144, 646–674 (2011). 10.1016/j.cell.2011.02.013

5 Seyfried, T. N. & Huysentruyt, L. C. On the origin of cancer metastasis. Crit Rev Oncog 18, 43–73 (2013).

6 Labelle, M. & Hynes, R. O. The initial hours of metastasis: the importance of cooperative host-tumor cell interactions during hematogenous dissemination. Cancer Discov 2, 1091–1099 (2012). 10.1158/2159-8290.CD-12-0329

7 Ridley, A. J. Rho GTPase signalling in cell migration. Curr Opin Cell Biol 36, 103–112 (2015). 10.1016/j.ceb.2015.08.005

8 Zhou, X. & Zheng, Y. Cell type-specific signaling function of RhoA GTPase: lessons from mouse gene targeting. J Biol Chem 288, 36179–36188 (2013). 10.1074/jbc.R113.515486

9 Thumkeo, D., Watanabe, S. & Narumiya, S. Physiological roles of Rho and Rho effectors in mammals. Eur J Cell Biol 92, 303–315 (2013). 10.1016/j.ejcb.2013.09.002

10 Jaffe, A. B. & Hall, A. Rho GTPases: biochemistry and biology. Annu Rev Cell Dev Biol 21, 247–269 (2005). 10.1146/annurev.cellbio.21.020604.150721

11 Iden, S. & Collard, J. G. Crosstalk between small GTPases and polarity proteins in cell polarization. Nat Rev Mol Cell Biol 9, 846–859 (2008). 10.1038/nrm2521

12 Burridge, K. Focal adhesions: a personal perspective on a half century of progress. FEBS J 284, 3355–3361 (2017). 10.1111/febs.14195

13 Heasman, S. J. & Ridley, A. J. Mammalian Rho GTPases: new insights into their functions from in vivo studies. Nat Rev Mol Cell Biol 9, 690–701 (2008). 10.1038/nrm2476

14 Strassheim, D. et al. RhoGTPase in Vascular Disease. Cells 8 (2019). 10.3390/cells8060551

15 Gavard, J. & Gutkind, J. S. Protein kinase C-related kinase and ROCK are required for thrombin-induced endothelial cell permeability downstream from Galpha12/13 and Galpha11/q. J Biol Chem 283, 29888–29896 (2008). 10.1074/jbc.M803880200

16 Basile, J. R., Gavard, J. & Gutkind, J. S. Plexin-B1 utilizes RhoA and Rho kinase to promote the integrin-dependent activation of Akt and ERK and endothelial cell motility. J Biol Chem 282, 34888–34895 (2007). 10.1074/jbc.M705467200

17 Mikelis, C. M. et al. RhoA and ROCK mediate histamine-induced vascular leakage and anaphylactic shock. Nat Commun 6, 6725 (2015). 10.1038/ncomms7725

18 Vandenbroucke, E., Mehta, D., Minshall, R. & Malik, A. B. Regulation of endothelial junctional permeability. Ann N Y Acad Sci 1123, 134–145 (2008). 10.1196/annals.1420.016

19 Carbajal, J. M. & Schaeffer, R. C., Jr. RhoA inactivation enhances endothelial barrier function. Am J Physiol 277, C955–964 (1999). 10.1152/ajpcell.1999.277.5.C955

20 Matsui, T. et al. Rho-associated kinase, a novel serine/threonine kinase, as a putative target for small GTP binding protein Rho. EMBO J 15, 2208–2216 (1996).

21 Amano, M. et al. Phosphorylation and activation of myosin by Rho-associated kinase (Rho-kinase). J Biol Chem 271, 20246–20249 (1996). 10.1074/jbc.271.34.20246

22 Ridley, A. J. & Hall, A. The small GTP-binding protein rho regulates the assembly of focal adhesions and actin stress fibers in response to growth factors. Cell 70, 389–399 (1992). 10.1016/0092-8674(92)90163-7

23 Chrzanowska-Wodnicka, M. & Burridge, K. Rho-stimulated contractility drives the formation of stress fibers and focal adhesions. J Cell Biol 133, 1403–1415 (1996). 10.1083/jcb.133.6.1403

24 Breslin, J. W. & Yuan, S. Y. Involvement of RhoA and Rho kinase in neutrophil-stimulated endothelial hyperpermeability. American journal of physiology. Heart and circulatory physiology 286, H1057–1062 (2004). 10.1152/ajpheart.00841.2003

25 van Nieuw Amerongen, G. P. & van Hinsbergh, V. W. Targets for pharmacological intervention of endothelial hyperpermeability and barrier function. Vascul Pharmacol 39, 257–272 (2002). 10.1016/s1537-1891(03)00014-4

26 Yao, L. et al. The role of RhoA/Rho kinase pathway in endothelial dysfunction. J Cardiovasc Dis Res 1, 165–170 (2010). 10.4103/0975-3583.74258

27 Huveneers, S., Daemen, M. J. & Hordijk, P. L. Between Rho(k) and a hard place: the relation between vessel wall stiffness, endothelial contractility, and cardiovascular disease. Circ Res 116, 895–908 (2015). 10.1161/CIRCRESAHA.116.305720

28 Gavard, J., Patel, V. & Gutkind, J. S. Angiopoietin-1 prevents VEGF-induced endothelial permeability by sequestering Src through mDia. Dev Cell 14, 25–36 (2008). 10.1016/j.devcel.2007.10.019

29 Zahra, F. T. et al. Endothelial RhoA GTPase is essential for in vitro endothelial functions but dispensable for physiological in vivo angiogenesis. Sci Rep 9, 11666 (2019). 10.1038/s41598-019-48053-z

30 Melendez, J. et al. RhoA GTPase is dispensable for actomyosin regulation but is essential for mitosis in primary mouse embryonic fibroblasts. J Biol Chem 286, 15132–15137 (2011). 10.1074/jbc.C111.229336

31 Ren, X. D., Kiosses, W. B. & Schwartz, M. A. Regulation of the small GTP-binding protein Rho by cell adhesion and the cytoskeleton. EMBO J 18, 578–585 (1999). 10.1093/emboj/18.3.578

32 Guilluy, C., Dubash, A. D. & Garcia-Mata, R. Analysis of RhoA and Rho GEF activity in whole cells and the cell nucleus. Nat Protoc 6, 2050–2060 (2011). 10.1038/nprot.2011.411

33 Mikelis, C., Sfaelou, E., Koutsioumpa, M., Kieffer, N. & Papadimitriou, E. Integrin alpha(v)beta(3) is a pleiotrophin receptor required for pleiotrophin-induced endothelial cell migration through receptor protein tyrosine phosphatase beta/zeta. FASEB J 23, 1459–1469 (2009). 10.1096/fj.08-117564

34 Koontz, L. TCA precipitation. Methods Enzymol 541, 3–10 (2014). 10.1016/B978-0-12-420119-4.00001-X

35 Mikelis, C. M. et al. PDZ-RhoGEF and LARG are essential for embryonic development and provide a link between thrombin and LPA receptors and Rho activation. J Biol Chem 288, 12232–12243 (2013). 10.1074/jbc.M112.428599

36 Sasportas, L. S., Hori, S. S., Pratx, G. & Gambhir, S. S. Detection and quantitation of circulating tumor cell dynamics by bioluminescence imaging in an orthotopic mammary carcinoma model. PLoS One 9, e105079 (2014). 10.1371/journal.pone.0105079

37 Rafehi, H. et al. Clonogenic assay: adherent cells. J Vis Exp (2011). 10.3791/2573

38 Gyorffy, B. et al. An online survival analysis tool to rapidly assess the effect of 22,277 genes on breast cancer prognosis using microarray data of 1,809 patients. Breast Cancer Res Treat 123, 725–731 (2010). 10.1007/s10549-009-0674-9

39 Bockhorn, M., Jain, R. K. & Munn, L. L. Active versus passive mechanisms in metastasis: do cancer cells crawl into vessels, or are they pushed? Lancet Oncol 8, 444–448 (2007). 10.1016/S1470-2045(07)70140-7

40 Mierke, C. T. Physical break-down of the classical view on cancer cell invasion and metastasis. Eur J Cell Biol 92, 89–104 (2013). 10.1016/j.ejcb.2012.12.002

41 Zervantonakis, I. K. et al. Three-dimensional microfluidic model for tumor cell intravasation and endothelial barrier function. Proc Natl Acad Sci U S A 109, 13515–13520 (2012). 10.1073/pnas.1210182109

42 Shenoy, A. K. & Lu, J. Cancer cells remodel themselves and vasculature to overcome the endothelial barrier. Cancer Lett 380, 534–544 (2016). 10.1016/j.canlet.2014.10.031

43 Srinivasan, B. et al. TEER measurement techniques for in vitro barrier model systems. J Lab Autom 20, 107–126 (2015). 10.1177/2211068214561025

44 Zomer, A. et al. In Vivo imaging reveals extracellular vesicle-mediated phenocopying of metastatic behavior. Cell 161, 1046–1057 (2015). 10.1016/j.cell.2015.04.042

45 Paltridge, J. L., Belle, L. & Khew-Goodall, Y. The secretome in cancer progression. Biochim Biophys Acta 1834, 2233–2241 (2013). 10.1016/j.bbapap.2013.03.014

46 Youngs, S. J., Ali, S. A., Taub, D. D. & Rees, R. C. Chemokines induce migrational responses in human breast carcinoma cell lines. Int J Cancer 71, 257–266 (1997). 10.1002/(sici)1097-0215(19970410)71:2<257::aid-ijc22>3.0.co;2-d

47 Dwyer, J. et al. Glioblastoma cell-secreted interleukin-8 induces brain endothelial cell permeability via CXCR2. PLoS One 7, e45562 (2012). 10.1371/journal.pone.0045562

48 Schraufstatter, I. U., Chung, J. & Burger, M. IL-8 activates endothelial cell CXCR1 and CXCR2 through Rho and Rac signaling pathways. Am J Physiol Lung Cell Mol Physiol 280, L1094–1103 (2001). 10.1152/ajplung.2001.280.6.L1094

49 Penna, G. et al. The vitamin D receptor agonist elocalcitol inhibits IL-8-dependent benign prostatic hyperplasia stromal cell proliferation and inflammatory response by targeting the RhoA/Rho kinase and NF-kappaB pathways. Prostate 69, 480–493 (2009). 10.1002/pros.20896

50 Lai, Y. et al. Interleukin-8 induces the endothelial cell migration through the activation of phosphoinositide 3-kinase-Rac1/RhoA pathway. Int J Biol Sci 7, 782–791 (2011). 10.7150/ijbs.7.782

51 Petreaca, M. L., Yao, M., Liu, Y., Defea, K. & Martins-Green, M. Transactivation of vascular endothelial growth factor receptor-2 by interleukin-8 (IL-8/CXCL8) is required for IL-8/CXCL8-induced endothelial permeability. Mol Biol Cell 18, 5014–5023 (2007). 10.1091/mbc.e07-01-0004

52 Lu, X. et al. Effective combinatorial immunotherapy for castration-resistant prostate cancer. Nature 543, 728–732 (2017). 10.1038/nature21676

53 Liao, W. et al. KRAS-IRF2 Axis Drives Immune Suppression and Immune Therapy Resistance in Colorectal Cancer. Cancer Cell 35, 559–572 e557 (2019). 10.1016/j.ccell.2019.02.008

54 Lawson, C. D. & Ridley, A. J. Rho GTPase signaling complexes in cell migration and invasion. J Cell Biol 217, 447–457 (2018). 10.1083/jcb.201612069

55 Kakiashvili, E. et al. GEF-H1 mediates tumor necrosis factor-alpha-induced Rho activation and myosin phosphorylation: role in the regulation of tubular paracellular permeability. J Biol Chem 284, 11454–11466 (2009). 10.1074/jbc.M805933200

56 Birukova, A. A. et al. Mechanotransduction by GEF-H1 as a novel mechanism of ventilator-induced vascular endothelial permeability. Am J Physiol Lung Cell Mol Physiol 298, L837–848 (2010). 10.1152/ajplung.00263.2009

57 Juettner, V. V. et al. VE-PTP stabilizes VE-cadherin junctions and the endothelial barrier via a phosphatase-independent mechanism. J Cell Biol 218, 1725–1742 (2019). 10.1083/jcb.201807210

58 Peng, J., He, F., Zhang, C., Deng, X. & Yin, F. Protein kinase C-alpha signals P115RhoGEF phosphorylation and RhoA activation in TNF-alpha-induced mouse brain microvascular endothelial cell barrier dysfunction. J Neuroinflammation 8, 28 (2011). 10.1186/1742-2094-8-28

59 Heemskerk, N. et al. F-actin-rich contractile endothelial pores prevent vascular leakage during leukocyte diapedesis through local RhoA signalling. Nat Commun 7, 10493 (2016). 10.1038/ncomms10493

60 Shang, X. et al. Small-molecule inhibitors targeting G-protein-coupled Rho guanine nucleotide exchange factors. Proc Natl Acad Sci U S A 110, 3155–3160 (2013). 10.1073/pnas.1212324110

61 Benoy, I. H. et al. Increased serum interleukin-8 in patients with early and metastatic breast cancer correlates with early dissemination and survival. Clin Cancer Res 10, 7157–7162 (2004). 10.1158/1078-0432.CCR-04-0812

62 Snoussi, K. et al. Genetic variation in IL-8 associated with increased risk and poor prognosis of breast carcinoma. Hum Immunol 67, 13–21 (2006). 10.1016/j.humimm.2006.03.018

63 Azevedo, A. S., Follain, G., Patthabhiraman, S., Harlepp, S. & Goetz, J. G. Metastasis of circulating tumor cells: favorable soil or suitable biomechanics, or both? Cell Adh Migr 9, 345–356 (2015). 10.1080/19336918.2015.1059563

64 Chambers, A. F., Groom, A. C. & MacDonald, I. C. Dissemination and growth of cancer cells in metastatic sites. Nat Rev Cancer 2, 563–572 (2002). 10.1038/nrc865

65 Follain, G. et al. Fluids and their mechanics in tumour transit: shaping metastasis. Nat Rev Cancer 20, 107–124 (2020). 10.1038/s41568-019-0221-x

66 Hamidi, H. & Ivaska, J. Every step of the way: integrins in cancer progression and metastasis. Nat Rev Cancer 18, 533–548 (2018). 10.1038/s41568-018-0038-z

67 Follain, G. et al. Hemodynamic Forces Tune the Arrest, Adhesion, and Extravasation of Circulating Tumor Cells. Dev Cell 45, 33–52 e12 (2018). 10.1016/j.devcel.2018.02.015

68 Reymond, N., d’Agua, B. B. & Ridley, A. J. Crossing the endothelial barrier during metastasis. Nat Rev Cancer 13, 858–870 (2013). 10.1038/nrc3628

69 Miles, F. L., Pruitt, F. L., van Golen, K. L. & Cooper, C. R. Stepping out of the flow: capillary extravasation in cancer metastasis. Clin Exp Metastasis 25, 305–324 (2008). 10.1007/s10585-007-9098-2

70 Lessey, E. C., Guilluy, C. & Burridge, K. From mechanical force to RhoA activation. Biochemistry 51, 7420–7432 (2012). 10.1021/bi300758e

71 Gould, R. A. et al. Cyclic Mechanical Loading Is Essential for Rac1-Mediated Elongation and Remodeling of the Embryonic Mitral Valve. Curr Biol 26, 27–37 (2016). 10.1016/j.cub.2015.11.033

72 Yamamoto, H. et al. Integrin beta1 controls VE-cadherin localization and blood vessel stability. Nat Commun 6, 6429 (2015). 10.1038/ncomms7429

73 Weis, S. M. & Cheresh, D. A. Tumor angiogenesis: molecular pathways and therapeutic targets. Nat Med 17, 1359–1370 (2011). 10.1038/nm.2537

74 Vogelsgesang, M., Pautsch, A. & Aktories, K. C3 exoenzymes, novel insights into structure and action of Rho-ADP-ribosylating toxins. Naunyn Schmiedebergs Arch Pharmacol 374, 347–360 (2007). 10.1007/s00210-006-0113-y

75 Muzumdar, M. D., Tasic, B., Miyamichi, K., Li, L. & Luo, L. A global double-fluorescent Cre reporter mouse. Genesis 45, 593–605 (2007). 10.1002/dvg.20335

76 Chen, M. T. et al. Comparison of patterns and prognosis among distant metastatic breast cancer patients by age groups: a SEER population-based analysis. Sci Rep 7, 9254 (2017). 10.1038/s41598-017-10166-8

77 Altorki, N. K. et al. The lung microenvironment: an important regulator of tumour growth and metastasis. Nat Rev Cancer 19, 9–31 (2019). 10.1038/s41568-018-0081-9

78 Minn, A. J. et al. Genes that mediate breast cancer metastasis to lung. Nature 436, 518–524 (2005). 10.1038/nature03799

79 Chauhan, V. P. et al. Reprogramming the microenvironment with tumor-selective angiotensin blockers enhances cancer immunotherapy. Proc Natl Acad Sci U S A 116, 10674–10680 (2019). 10.1073/pnas.1819889116

80 Hiratsuka, S. et al. Endothelial focal adhesion kinase mediates cancer cell homing to discrete regions of the lungs via E-selectin up-regulation. Proc Natl Acad Sci U S A 108, 3725–3730 (2011). 10.1073/pnas.1100446108

81 Dagogo-Jack, I. & Shaw, A. T. Tumour heterogeneity and resistance to cancer therapies. Nat Rev Clin Oncol 15, 81–94 (2018). 10.1038/nrclinonc.2017.166

82 Al-Husein, B., Abdalla, M., Trepte, M., Deremer, D. L. & Somanath, P. R. Antiangiogenic therapy for cancer: an update. Pharmacotherapy 32, 1095–1111 (2012). 10.1002/phar.1147

83 Hiratsuka, S. et al. MMP9 induction by vascular endothelial growth factor receptor-1 is involved in lung-specific metastasis. Cancer Cell 2, 289–300 (2002).

84 Kaplan, R. N. et al. VEGFR1-positive haematopoietic bone marrow progenitors initiate the pre-metastatic niche. Nature 438, 820–827 (2005). 10.1038/nature04186

85 Hiratsuka, S., Watanabe, A., Aburatani, H. & Maru, Y. Tumour-mediated upregulation of chemoattractants and recruitment of myeloid cells predetermines lung metastasis. Nat Cell Biol 8, 1369–1375 (2006). 10.1038/ncb1507

86 Khanna, C. & Hunter, K. Modeling metastasis in vivo. Carcinogenesis 26, 513–523 (2005). 10.1093/carcin/bgh261

87 Gomez-Cuadrado, L., Tracey, N., Ma, R., Qian, B. & Brunton, V. G. Mouse models of metastasis: progress and prospects. Dis Model Mech 10, 1061–1074 (2017). 10.1242/dmm.030403

88 van Nieuw Amerongen, G. P., van Delft, S., Vermeer, M. A., Collard, J. G. & van Hinsbergh, V. W. Activation of RhoA by thrombin in endothelial hyperpermeability: role of Rho kinase and protein tyrosine kinases. Circ Res 87, 335–340 (2000). 10.1161/01.res.87.4.335

89 Maekawa, M. et al. Signaling from Rho to the actin cytoskeleton through protein kinases ROCK and LIM-kinase. Science 285, 895–898 (1999). 10.1126/science.285.5429.895

90 McKenzie, J. A. & Ridley, A. J. Roles of Rho/ROCK and MLCK in TNF-alpha-induced changes in endothelial morphology and permeability. J Cell Physiol 213, 221–228 (2007). 10.1002/jcp.21114

91 Sun, H., Breslin, J. W., Zhu, J., Yuan, S. Y. & Wu, M. H. Rho and ROCK signaling in VEGF-induced microvascular endothelial hyperpermeability. Microcirculation 13, 237–247 (2006). 10.1080/10739680600556944

92 Ma, T., Liu, L., Wang, P. & Xue, Y. Evidence for involvement of ROCK signaling in bradykinin-induced increase in murine blood-tumor barrier permeability. J Neurooncol 106, 291–301 (2012). 10.1007/s11060-011-0685-3

93 Uehata, M. et al. Calcium sensitization of smooth muscle mediated by a Rho-associated protein kinase in hypertension. Nature 389, 990–994 (1997). 10.1038/40187

94 Asano, T. et al. Mechanism of action of a novel antivasospasm drug, HA1077. J Pharmacol Exp Ther 241, 1033–1040 (1987).

95 Wei, L., Surma, M., Shi, S., Lambert-Cheatham, N. & Shi, J. Novel Insights into the Roles of Rho Kinase in Cancer. Arch Immunol Ther Exp (Warsz) 64, 259–278 (2016). 10.1007/s00005-015-0382-6

96 Vennin, C. et al. Transient tissue priming via ROCK inhibition uncouples pancreatic cancer progression, sensitivity to chemotherapy, and metastasis. Sci Transl Med 9 (2017). 10.1126/scitranslmed.aai8504

97 Ying, H. et al. The Rho kinase inhibitor fasudil inhibits tumor progression in human and rat tumor models. Mol Cancer Ther 5, 2158–2164 (2006). 10.1158/1535-7163.MCT-05-0440

98 Guerra, F. S., Oliveira, R. G., Fraga, C. A. M., Mermelstein, C. D. S. & Fernandes, P. D. ROCK inhibition with Fasudil induces beta-catenin nuclear translocation and inhibits cell migration of MDA-MB 231 human breast cancer cells. Sci Rep 7, 13723 (2017). 10.1038/s41598-017-14216-z

99 Tachibana, E. et al. Intra-arterial infusion of fasudil hydrochloride for treating vasospasm following subarachnoid haemorrhage. Acta Neurochir (Wien) 141, 13–19 (1999). 10.1007/s007010050260

100 Shibuya, M. et al. Effects of fasudil in acute ischemic stroke: results of a prospective placebo-controlled double-blind trial. J Neurol Sci 238, 31–39 (2005). 10.1016/j.jns.2005.06.003

101 He, B. et al. Remodeling of Metastatic Vasculature Reduces Lung Colonization and Sensitizes Overt Metastases to Immunotherapy. Cell Rep 30, 714–724 e715 (2020). 10.1016/j.celrep.2019.12.013

102 Gilbert-Ross, M., Marcus, A. I. & Zhou, W. RhoA, a novel tumor suppressor or oncogene as a therapeutic target? Genes Dis 2, 2–3 (2015). 10.1016/j.gendis.2014.10.001

103 O’Hayre, M. et al. Inactivating mutations in GNA13 and RHOA in Burkitt’s lymphoma and diffuse large B-cell lymphoma: a tumor suppressor function for the Galpha13/RhoA axis in B cells. Oncogene 35, 3771–3780 (2016). 10.1038/onc.2015.442

104 Barlow, H. R. & Cleaver, O. Building Blood Vessels-One Rho GTPase at a Time. Cells 8 (2019). 10.3390/cells8060545

105 Radeva, M. Y. & Waschke, J. Mind the gap: mechanisms regulating the endothelial barrier. Acta Physiol (Oxf) 222 (2018). 10.1111/apha.12860

106 Bauer, K., Mierke, C. & Behrens, J. Expression profiling reveals genes associated with transendothelial migration of tumor cells: a functional role for alphavbeta3 integrin. Int J Cancer 121, 1910–1918 (2007). 10.1002/ijc.22879

107 Gay, L. J. & Felding-Habermann, B. Contribution of platelets to tumour metastasis. Nat Rev Cancer 11, 123–134 (2011). 10.1038/nrc3004

108 Kopf, A. et al. Microtubules control cellular shape and coherence in amoeboid migrating cells. J Cell Biol 219 (2020). 10.1083/jcb.201907154

109 Lahooti, B. et al. Endothelial-Specific Targeting of RhoA Signaling via CD31 Antibody-Conjugated Nanoparticles. J Pharmacol Exp Ther 385, 35–49 (2023). 10.1124/jpet.122.001384

110 Mierke, C. T., Frey, B., Fellner, M., Herrmann, M. & Fabry, B. Integrin alpha5beta1 facilitates cancer cell invasion through enhanced contractile forces. J Cell Sci 124, 369–383 (2011). 10.1242/jcs.071985

111 Stoletov, K. et al. Visualizing extravasation dynamics of metastatic tumor cells. J Cell Sci 123, 2332–2341 (2010). 10.1242/jcs.069443

112 Kim, J. et al. Topological Adaptation of Transmembrane Domains to the Force-Modulated Lipid Bilayer Is a Basis of Sensing Mechanical Force. Curr Biol 30, 1614–1625 e1615 (2020). 10.1016/j.cub.2020.02.028

113 Paszek, M. J. et al. Tensional homeostasis and the malignant phenotype. Cancer Cell 8, 241–254 (2005). 10.1016/j.ccr.2005.08.010

114 Acerbi, I. et al. Human breast cancer invasion and aggression correlates with ECM stiffening and immune cell infiltration. Integr Biol (Camb) 7, 1120–1134 (2015). 10.1039/c5ib00040h

115 Hakanpaa, L. et al. Endothelial destabilization by angiopoietin-2 via integrin beta1 activation. Nat Commun 6, 5962 (2015). 10.1038/ncomms6962

116 Akwii, R. G. et al. Angiopoietin-2-induced lymphatic endothelial cell migration drives lymphangiogenesis via the beta1 integrin-RhoA-formin axis. Angiogenesis 25, 373–396 (2022). 10.1007/s10456-022-09831-y

117 Shao, S. et al. Targeting High Expressed alpha5beta1 Integrin in Liver Metastatic Lesions To Resist Metastasis of Colorectal Cancer by RPM Peptide-Modified Chitosan-Stearic Micelles. Mol Pharm 15, 1653–1663 (2018). 10.1021/acs.molpharmaceut.8b00013

118 Sun, Z. et al. Single-cell RNA sequencing reveals gene expression signatures of breast cancer-associated endothelial cells. Oncotarget 9, 10945–10961 (2018). 10.18632/oncotarget.23760

119 Collo, G. & Pepper, M. S. Endothelial cell integrin alpha5beta1 expression is modulated by cytokines and during migration in vitro. J Cell Sci 112 (Pt 4), 569–578 (1999).

120 Reymond, N. et al. Cdc42 promotes transendothelial migration of cancer cells through beta1 integrin. J Cell Biol 199, 653–668 (2012). 10.1083/jcb.201205169

121 Joyce, J. A. & Pollard, J. W. Microenvironmental regulation of metastasis. Nat Rev Cancer 9, 239–252 (2009). 10.1038/nrc2618

122 Haga, R. B. & Ridley, A. J. Rho GTPases: Regulation and roles in cancer cell biology. Small GTPases 7, 207–221 (2016). 10.1080/21541248.2016.1232583

123 Orgaz, J. L. et al. Myosin II Reactivation and Cytoskeletal Remodeling as a Hallmark and a Vulnerability in Melanoma Therapy Resistance. Cancer Cell 37, 85–103 e109 (2020). 10.1016/j.ccell.2019.12.003

124 Feng, Y., LoGrasso, P. V., Defert, O. & Li, R. Rho Kinase (ROCK) Inhibitors and Their Therapeutic Potential. J Med Chem 59, 2269–2300 (2016). 10.1021/acs.jmedchem.5b00683

125 Ethier, S. P., Kokeny, K. E., Ridings, J. W. & Dilts, C. A. erbB family receptor expression and growth regulation in a newly isolated human breast cancer cell line. Cancer Res 56, 899–907 (1996).

126 Ewens, A., Mihich, E. & Ehrke, M. J. Distant metastasis from subcutaneously grown E0771 medullary breast adenocarcinoma. Anticancer Res 25, 3905–3915 (2005).

127 Wang, Y. et al. Ephrin-B2 controls VEGF-induced angiogenesis and lymphangiogenesis. Nature 465, 483–486 (2010). 10.1038/nature09002

