## Supplemental Figures, Figure legends and text for "Tumor-induced endothelial RhoA activation mediates tumor cell transendothelial migration and metastasis"

#### Supplemental Figure Legends

**Supplemental Figure 1.**

(A) Schematic diagram of in vitro transendothelial migration and transendothelial electrical resistance (TEER) measurement, with cancer (MDA-MB-231) (i), endothelial (EAhy926) (ii) or no cells (Medium) (iii) on the upper surface of the transwell membrane.

(B) Measurement of TEER across endothelial monolayer before and after the addition of $2x10^5$ endothelial (EAhy926) or cancer (MDA-MB-231) cells or serum-free medium (control). Dotted box designates the increase of TEER following cell or medium addition n = 2.

(C) Designation of each cytokine presented in the graph in figure 1I.

**Supplemental Figure 2.**

(A and B) Representative images of RhoA activation in HUVECs upon (A) sTNF-R1 (5 ng/ml) and (B) BAFF (8 ng/ml) treatment at different time points. sTNF-R1, n = 3; BAFF, n = 4.

(C) Quantification of migrated SUM52PE tumor cells in the absence or presence of IL-8 (10 ng/ml) after overnight treatment with DMSO (control) or SX-682 (2 μM) (nuclei = blue). n = 4.

(D and E) Representative images (H) and quantification (I) of transendothelial migration of green fluorescent dye-labeled SUM52PE cancer cells through an EAhy926 monolayer on collagen-coated transwell inserts in absence or presence of recombinant IL-8 (10 ng/ml) after 24 h (nuclei = blue); n = 2. Scale bars, 100 μm.

(F) Quantification of IL-8 mRNA levels of siNeg- and siIL-8-transfected MDA-MB-231 cells at 48 h post-transfection; n = 2.

(G) Quantification of IL-8 protein levels of siNeg- and siIL-8-transfected MDA-MB-231 cells at 48 h post-transfection; n = 2.

(H and I) Representative images (H) and quantification (I) of siNeg- and siIL-8-transfected MDA-MB-231 cancer cell migration (nuclei = blue); n = 3. Scale bars, 100 μm.

(J and K) Representative images (J) and quantification (K) of RhoA activation in HUVECs upon treatment of conditioned medium from siNeg- or siIL-8- transfected MDA-MB-231 at 5 min; n = 2.

(L and M) Representative images (L) and quantification (M) of RhoA activation in HUVEC 5 min after treatment with IL-8 (10 ng/ml) in the absence or presence of the SX-682 (2 μM); n = 3.

(N) Quantification of IL-8-induced PDZ RhoGEF, LARG and GEF-H1 activation in HUVECs upon 5 min of IL-8 (10 ng/ml) incubation; PDZ RhoGEF, n = 2; LARG, n = 3; GEF-H1, n = 3.

(O and P) Representative images (O) and quantification (P) of p115-RhoGEF knockdown in EAhy926 by siRNA treatment (50 nM) at 48 h post-transfection; n = 2.

(Q and R) Representative images (Q) and quantification (R) of transendothelial migration of GFP-expressing MDA-MB-231 cancer cells across EAhy926 cells transfected with siNeg or sip115-RhoGEF. n = 4; Scale bars, 100 μm.

Data in (E), (F), (G), (I), (K), (P), (N) and (R) represent mean ± SEM and were analyzed by Student's unpaired t-test. Data in (C) and (M) were analyzed by one-way ANOVA. ns = not significant; *P < 0.05, **P < 0.01.

**Supplemental Figure 3.**

(A and B) Representative images (A) and quantification (B) of IL-8-induced p115RhoGEF activation in HUVECs upon 5 min of IL-8 (10 ng/ml) incubation with or without Y16 pretreatment; n = 4.

(C and D) Representative images (C) and quantification (D) of transendothelial migration of GFP-expressing MDA-MB-231 cancer cells across EAhy926 cells in absence or presence of IL-8 (10 ng/ml) with or without Y16 pretreatment. n = 2; Scale bars, 100 µm.

(E and F) Metastasis-free survival of lymph node negative breast cancer patients, stratified on expression of two IL-8 isoforms: 77 amino acids (E) and 72 amino acids (F) expression, respectively.

(G and H) Metastasis-free survival of breast cancer patients, stratified on sTNF-R1 (G) and BAFF (H) expression, respectively.

Data in (B) and (D) represent mean ± SEM and were analyzed by one-way ANOVA. Data in (E), (F), (G) and (H) were analyzed by Kaplan-Meier estimator. ns = not significant; *P < 0.05, **P < 0.01; ***P < 0.001.

**Supplemental Figure 4.**

(A) Representative images of cancer-cancer (CC/CC), endothelial-endothelial (EC/EC), or endothelial-cancer (EC/CC) contact points during co-culture of HUVEC and GFP+ MDA-MB-231 cancer cells.

(B-C) Representative images (B) and quantification (C) of RhoA activation at the cell-to-cell contact points during co-culture of HUVEC and green fluorescent dye-labeled BEAS-2B (epithelial) cells; n = 2.

(D) Representative images of pMLC and phalloidin staining during co-culture of HUVEC and GFP+ MDA-MB-231 cancer cells; Scale bars, 20 μm.

(E) Representative images of transendothelial migration of GFP+ MDA-MB-231 cells after endothelial pretreatment with control or RGD peptide; Scale bars, 100 μm.

(F) Representative images of transendothelial migration of GFP+ MDA-MB-231 cells after endothelial pretreatment with Isotype IgG or blocking antibodies against integrins $\alpha_5\beta_1$ and $\alpha_V\beta_3$; Scale bars, 100 μm.

(G) Representative images of active $\beta_1$ integrin staining during co-culture of HUVEC and GFP+ MDA-MB-231 cancer cells; Scale bars, 20 μm.

(H and I) Quantification of integrin $\beta_1$ (H) and $\beta_3$ (I) knockdown in EAhy926 cells by siRNA treatment (50 nM) at 48 h post-transfection; n = 4 and n = 3 respectively.

(J) Representative images of transendothelial migration of GFP+ MDA-MB-231 cells after treatment of the endothelial cells with scramble siRNA (Neg) or siRNA for integrins $\beta_1$ or $\beta_3$; Scale bars, 100 μm.

(K) Representative images of phalloidin fluorescence intensity at the cell-to-cell contact points during co-culture of GFP+ MDA-MB-231 cancer cells and HUVEC treated with scramble siRNA (Neg) or siRNA for integrins $\beta_1$ or $\beta_3$; Scale bars, 10 μm.

(L) Representative images of transendothelial migration of GFP+ MDA-MB-231 cells after treatment of the endothelial cells with the Src inhibitor (PP2) and the corresponding control (PP3); Scale bars, 100 μm.

(M) Representative images of transendothelial migration of GFP+ MDA-MB-231 cells after treatment of the endothelial cells with vehicle (Control) or the Src inhibitor (SU6656); Scale bars, 100 μm.

Data in (C), (H) and (I) represent mean ± SEM. Data in (C) were analyzed by one-way ANOVA and data in (H) and (I) and were analyzed by Student’s unpaired t-test. ns = not significant; ***$P < 0.001$.

**Supplemental Figure 5.**

(A) Representative images of RhoA pulldown experiments on HUVEC treated with vehicle, RGD peptide and blocking antibody against $\alpha_5\beta_1$ in the presence or absence of IL-8 treatment; n = 3.

(B-D) Evaluation of metastatic outcome of B16 syngeneic melanoma cancer cell line (I.V.) in mice following treatment with β1 blocking antibody and SX-682. (B) Representative IVIS images of the treated animals. (C) Representative macroscopic appearance of the dissected lungs. (D) Quantification of number of metastases per mouse; Control, n = 10; Treatment, n = 10.

Data in (D) represent mean ± SEM and was analyzed by Student's unpaired t-test. ***P < 0.001.

**Supplemental figure 6.**

(A-C) Representative images and quantification of RhoA activation in HUVECs upon incubation with serum-free conditioned medium from EO771 (A), B16-F10 (B) and LLC (C) cancer cells; A, n = 4; B, n = 3; C, n = 4.

(D and E) Representative images (D) and quantification (E) of trans-endothelial migration of GFP-labeled MDA-MB-231 cancer cells in the absence or presence of IL-8 (50 ng/ml) through EAhy926 endothelial monolayer with or without C3 toxin pretreatment (nuclei = blue); n = 4. Scale bars, 100 μm.

(F and G) Representative images (F) and quantification (G) of trans-endothelial migration of GFP-labeled NIH-3T3 cells through EAhy926 endothelial monolayer with or without C3 toxin pretreatment (nuclei = blue); n = 3. Scale bars, 100 μm. (H and I) Representative images (H) and quantification (I) of RhoA knockdown in EAhy926 cells by siRNA treatment (50 nM) at 48 h post-transfection; n = 3.

(J and K) Representative images (J) and quantification (K) of trans-endothelial migration of GFP+ MDA-MB-231 cancer cells through siNeg- or siRhoA-transfected (50 nM) EAhy926 endothelial monolayer (nuclei = blue); n = 3. Scale bars, 100 μm.

Data in (A), (B), (C), (E), (G), (I), (K) represent mean ± SEM. Data in (A), (B), (C), (E) and (K) was analyzed by one way ANOVA and data in (G) and (I) was analyzed by Student’s unpaired t test. ns = not significant; *P < 0.05; **P < 0.01; ***P < 0.001.

**Supplemental figure 7.**

(A-D) Evaluation of metastatic outcome of B16-F10 syngeneic melanoma cell line in $RhoA^{i\Delta EC}$ mice and littermate controls (IC administration). (A) Representative IVIS images of the mice and dissected lungs. (B) Representative macroscopic appearance of the dissected lungs (arrows denote metastases). (C) Quantification of number of metastases per mouse; Control, n = 12; $RhoA^{i\Delta EC}$, n = 12. (D) Representative H&E lung sections. Scale bars, 1 mm.

(E-H) Evaluation of metastatic outcome of LLC syngeneic lung cancer cell line in $RhoA^{i\Delta EC}$ mice and littermate controls (IC administration). (E) Representative IVIS images of the mice and dissected lungs. (F) Representative macroscopic appearance of the dissected lungs (arrows denote metastases). (G) Quantification of number of metastases per mouse; Control, n = 11; $RhoA^{i\Delta EC}$, n = 12. (H) Representative H&E lung sections. Scale bars, 1 mm.

(I-L) Evaluation of metastatic outcome of EO771 syngeneic breast cancer cells in $RhoA^{i\Delta EC}$ mice and littermate controls (orthotopic breast cancer spontaneous metastasis model). (I) Representative IVIS images of the mice. (J) Quantification of number of metastases per mouse. (K) Representative IVIS images of colonies from circulating tumor cells isolated from blood. (L) Quantification of number of clonies from each mice; Control, n = 6; $RhoA^{i\Delta EC}$, n = 5. Data in (C), (G), (J) and (L) represent mean ± SEM and were analyzed by Student's unpaired t-test. *P < 0.05; ***P < 0.001.

**Supplemental figure 8.**

(A-D) Evaluation of metastatic outcome of B16-F10 syngeneic melanoma cell line (I.C.) in mice treated daily with vehicle or Fasudil (20 mg/kg; I.P.). (A) Representative IVIS images of the mice and dissected lungs. (B) Representative macroscopic appearance of the dissected lungs. (C) Quantification of number of metastases per mouse; Control, n = 5; Fasudil, n = 4. (D) Representative H&E lung sections. Scale bars, 1 mm.

(E-H) Evaluation of metastatic outcome of LLC syngeneic lung cancer cell line (I.C.) in mice treated daily with vehicle or Fasudil (20 mg/kg; I.P.). (E) Representative IVIS images of the mice and dissected lungs. (F) Representative macroscopic appearance of the dissected lungs (arrows denote metastases). (G) Quantification of number of metastases per mouse; Control, n = 10; Fasudil, n = 8. (H) Representative H&E lung sections. Scale bars, 1 mm.

(I-K) Evaluation of metastatic outcome of B16 syngeneic melanoma cell line (I.V.) in mice treated daily with vehicle or Fasudil, Fasudil before: treatment started before the administration of the cancer cells; Fasudil after: treatment started after the administration of the cancer cells (20 mg/kg; I.P.). (I) Representative IVIS images of the mice. (J) Representative macroscopic appearance of the dissected lungs. (K) Quantification of number of metastases per mouse; Control, n = 10; Fasudil before, n = 10; Fasudil before, n = 9.

(L and M) Evaluation of extravasation of B16-F10 syngeneic melanoma cells (I.V.) in mice treated with vehicle or Fasudil. Representative images (L) and quantification (M) of colonies grown from isolated lungs; Control, n = 6; Fasudil= 9.

Data in (C), (G), (K) and (M) represent mean ± SEM. Data in (C), (G) and (M) was analyzed by Student's unpaired t-test and data in (K) was analyzed by one-way ANOVA. *P < 0.05, ***P < 0.001.

**Supplemental figure 9. Fasudil administration does not present toxicity indications.**

(A-D) Quantification of mouse weight during intravenous experimental metastasis models with (A) EO771, (B) B16-F10, (C) LLC murine syngeneic and (D) MDA-MB-231 human tumor cells. (A) Control, n = 8; Fasudil, n = 7; (B) Control, n = 14; Fasudil, n = 14; (C) Control, n = 7; Fasudil, n = 9; (D) Control, n = 13; Fasudil, n = 14.

(E-F) Quantification of mouse weight during intracardiac experimental metastasis models with (E) B16-F10 and (F) LLC murine syngeneic cancer cells. (E) Control, n = 5; Fasudil, n = 4; (F) Control, n = 10; Fasudil, n = 8.

(G) Quantification of mouse weight for the experiment described in supplemental figure 6I-K; Control, n = 10; Fasudil before, n = 10; Fasudil before, n = 9.

(H) Quantification of mouse weight for the experiment described in supplemental figure 4N-P; Control, n = 10; Treatment, n = 10.

Data in (A), (B), (C), (D) (E), (F), (G) and (H) represent mean ± SEM.

**Parameters used to create the graphs presented in figures 2P**

Figure 2N was generated with the Kaplan Meier Plotter for Breast Cancer online tool (http://kmplot.com/analysis/index.php?p=service&cancer=breast)

Affy ID: 211506_s_at (IL8)

Survival: DMFS

Auto select best cutoff: checked

Follow up threshold: all

Censore at threshold: checked

Compute median over entire database: false

Cutoff value used in analysis: 65

Expression range of the probe: 1 - 6258

Probe set option: user selected probe set

Invert HR values below 1: not checked

**Restrictions**

ER status: all

derive ER status from gene expression data: not checked

PR status: all

HER2 status: all

Lymph node status: all

Intrinsic subtype: all

TP53 status: all

Pietenpol subtype: all

Grade: all

Use earlier release of the database: all

Use following dataset for the analysis: all

**Quality control**

Remove redundant samples: checked

Array quality control: exclude biased arrays

Proportional hazards assumption: 0

**Cohort**

Cohorts: not selected

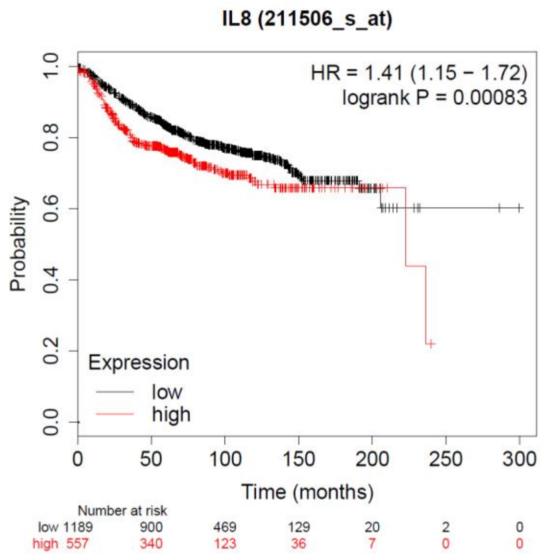

**Parameters used to create the graphs presented in figure 2Q**

Figure 2O was generated with the Kaplan Meier Plotter for Breast Cancer online tool (http://kmplot.com/analysis/index.php?p=service&cancer=breast)

Affy ID: 202859_x_at (MDNCF)

Survival: DMFS

Auto select best cutoff: checked

Follow up threshold: all

Censore at threshold: checked

Compute median over entire database: false

Cutoff value used in analysis: 286

Expression range of the probe: 5 - 19696

Probe set option: user selected probe set

Invert HR values below 1: not checked

**Restrictions**

ER status: all

derive ER status from gene expression data: not checked

PR status: all

HER2 status: all

Lymph node status: all

Intrinsic subtype: all

TP53 status: all

Pietenpol subtype: all

Grade: all

Use earlier release of the database: all

Use following dataset for the analysis: all

**Quality control**

Remove redundant samples: checked

Array quality control: exclude biased arrays

Proportional hazards assumption: 0

**Cohort**

Cohorts: not selected

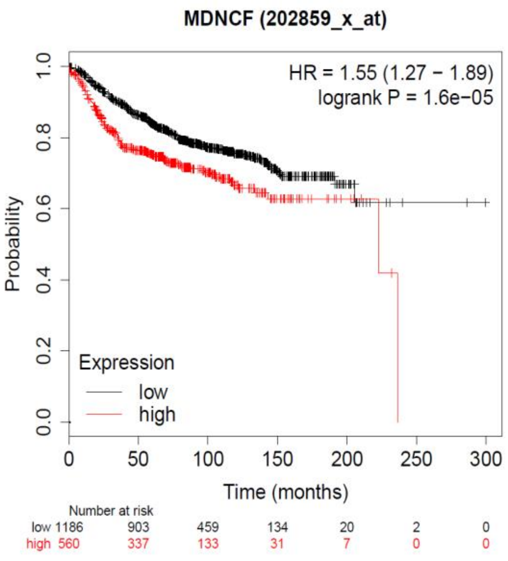

**Parameters used to create the graphs presented in supplemental figure 3E**

Supplemental figure 2J was generated with the Kaplan Meier Plotter for Breast Cancer online tool

(http://kmplot.com/analysis/index.php?p=service&cancer=breast)

Affy ID: 211506_s_at (IL8)

Survival: DMFS

Auto select best cutoff: checked

Follow up threshold: all

Censore at threshold: checked

Compute median over entire database: false

Cutoff value used in analysis: 75

Expression range of the probe: 1-6258

Probe set option: user selected probe set

Invert HR values below 1: not checked

**Restrictions**

ER status: all

derive ER status from gene expression data: not checked

PR status: all

HER2 status: all

Lymph node status: Lymph node negative

Intrinsic subtype: all

TP53 status: all

Pietenpol subtype: all

Grade: all

Use earlier release of the database: all

Use following dataset for the analysis: all

**Quality control**

Remove redundant samples: checked; Array quality control: exclude biased arrays; Proportional hazards assumption: 0

**Cohort**

Cohorts: not selected

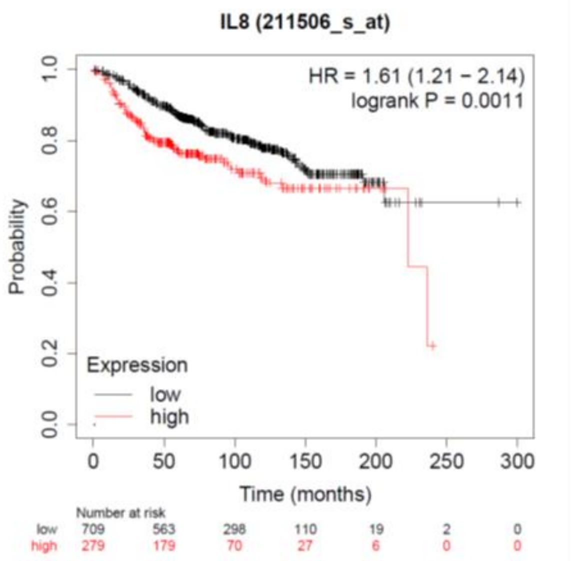

**Parameters used to create the graphs presented in supplemental figure 3F**

Supplemental figure 2K was generated with the Kaplan Meier Plotter for Breast Cancer online tool

(http://kmplot.com/analysis/index.php?p=service&cancer=breast)

Affy ID: 202859_x_at (MDNCF)

Survival: DMFS

Auto select best cutoff: checked

Follow up threshold: all

Censore at threshold: checked

Compute median over entire database: false

Cutoff value used in analysis: 203

Expression range of the probe: 5-6258

Probe set option: user selected probe set

Invert HR values below 1: not checked

**Restrictions**

ER status: all

derive ER status from gene expression data: not checked

PR status: all; HER2 status: all

Lymph node status: Lymph node negative

Intrinsic subtype: all

TP53 status: all

Pietenpol subtype: all

Grade: all; Use earlier release of the database: all

Use following dataset for the analysis: all

**Quality control**

Remove redundant samples: checked; Array quality control: exclude biased arrays; Proportional hazards assumption: 0

**Cohort**

Cohorts: not selected

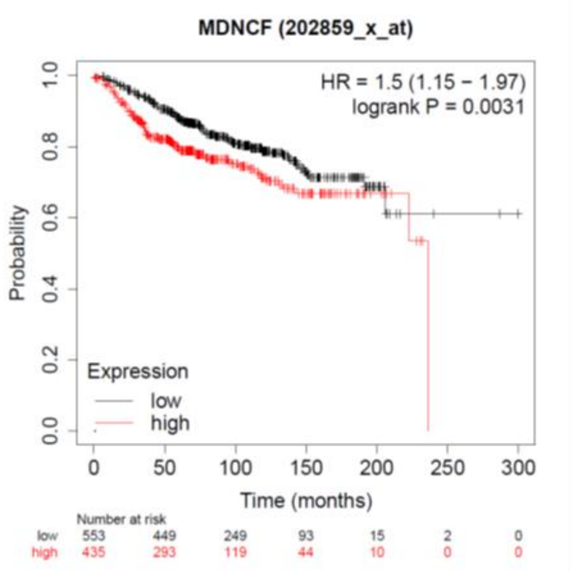

**Parameters used to create the graphs presented in supplemental figure 3G**

Supplemental figure 2L was generated with the Kaplan Meier Plotter for Breast Cancer online tool

(http://kmplot.com/analysis/index.php?p=service&cancer=breast)

Affy ID: 207643_s_at (TNF−R)

Survival: DMFS

Auto select best cutoff: checked

Follow up threshold: all

Censore at threshold: checked

Compute median over entire database: false

Cutoff value used in analysis: 1170

Expression range of the probe: 221-11087

Probe set option: user selected probe set

Invert HR values below 1: not checked

**Restrictions**

ER status: all

derive ER status from gene expression data: not checked

PR status: all

HER2 status: all

Lymph node status: all

Intrinsic subtype: all

TP53 status: all

Pietenpol subtype: all

Grade: all

Use earlier release of the database: all

Use following dataset for the analysis: all

**Quality control**

Remove redundant samples: checked

Array quality control: exclude biased arrays

Proportional hazards assumption: 0

**Cohort**

Cohorts: not selected

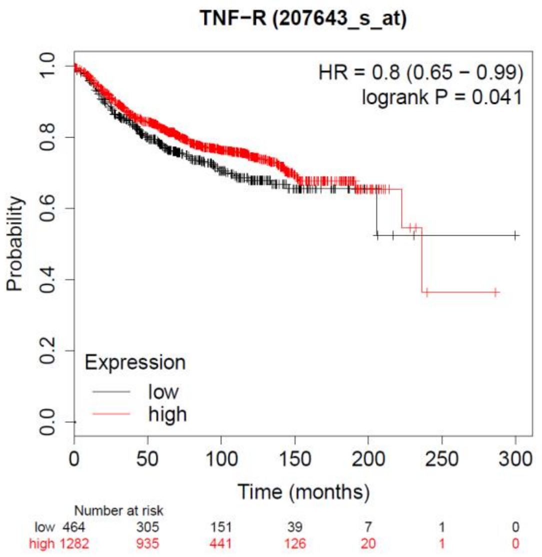

**Parameters used to create the graphs presented in supplemental figure 3H**

Supplemental figure 2M was generated with the Kaplan Meier Plotter for Breast Cancer online tool

(http://kmplot.com/analysis/index.php?p=service&cancer=breast)

Affy ID: 207641_at (TNFRSF13B/BAFF)

Survival: DMFS

Auto select best cutoff: checked

Follow up threshold: all

Censore at threshold: checked

Compute median over entire database: false

Cutoff value used in analysis: 103

Expression range of the probe: 9 - 2503

Probe set option: user selected probe set

Invert HR values below 1: not checked

**Restrictions**

ER status: all

derive ER status from gene expression data: not checked

PR status: all

HER2 status: all

Lymph node status: all

Intrinsic subtype: all

TP53 status: all

Pietenpol subtype: all

Grade: all

Use earlier release of the database: all

Use following dataset for the analysis: all

**Quality control**

Remove redundant samples: checked

Array quality control: exclude biased arrays

Proportional hazards assumption: 0

**Cohort**

Cohorts: not selected

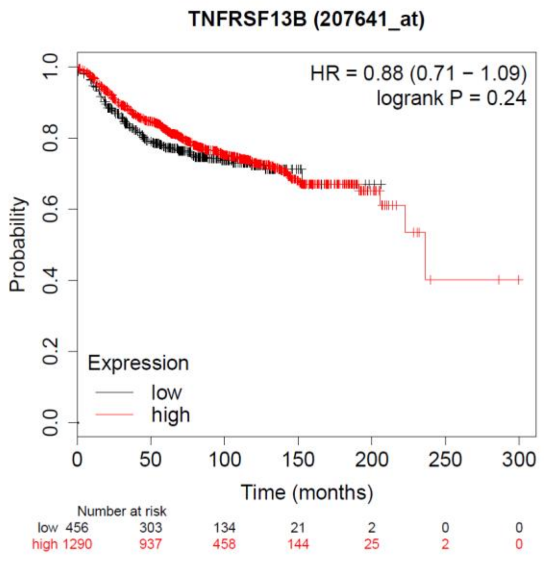

**Supplemental Figure 1.**

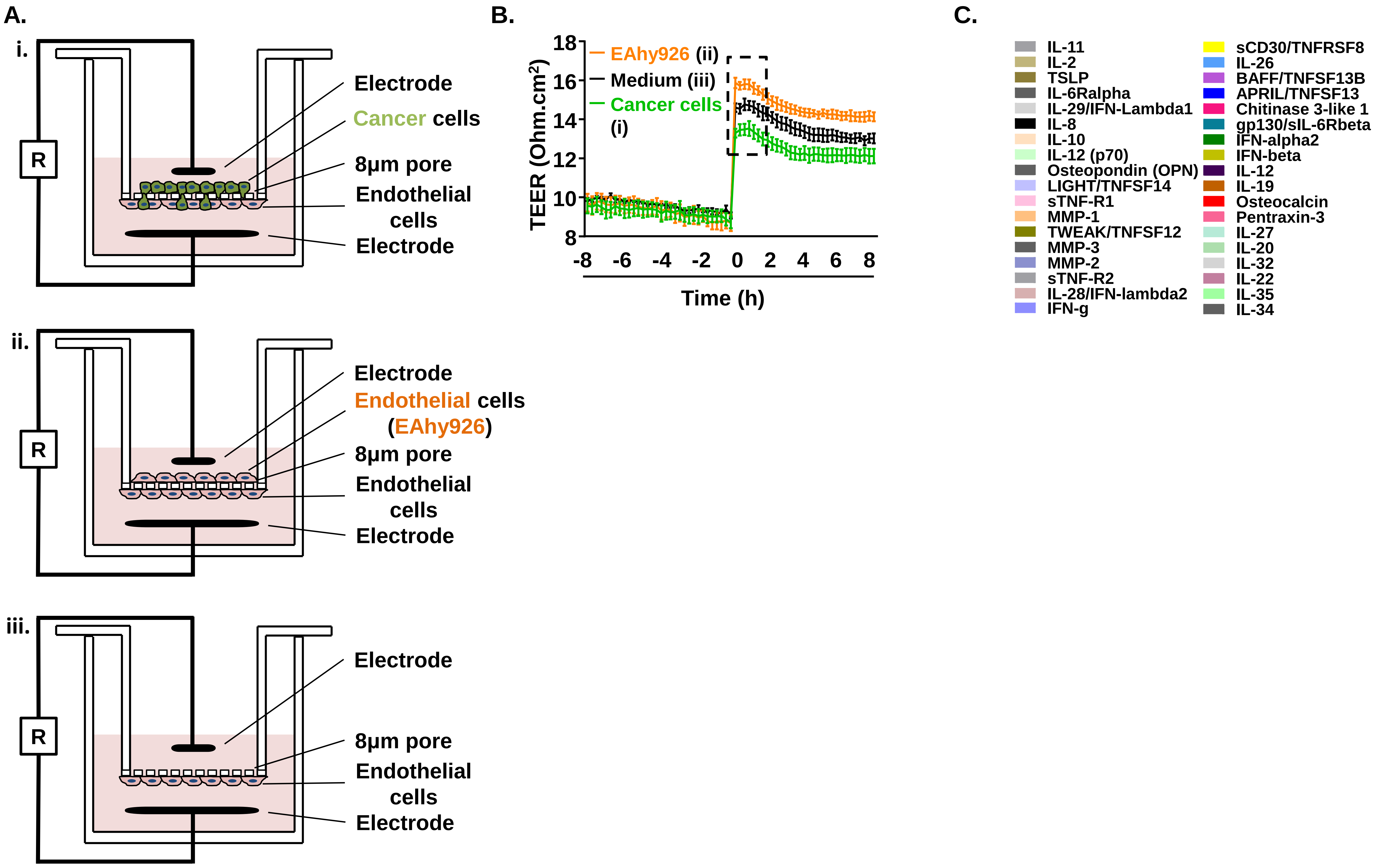

**Supplemental Figure 2.**

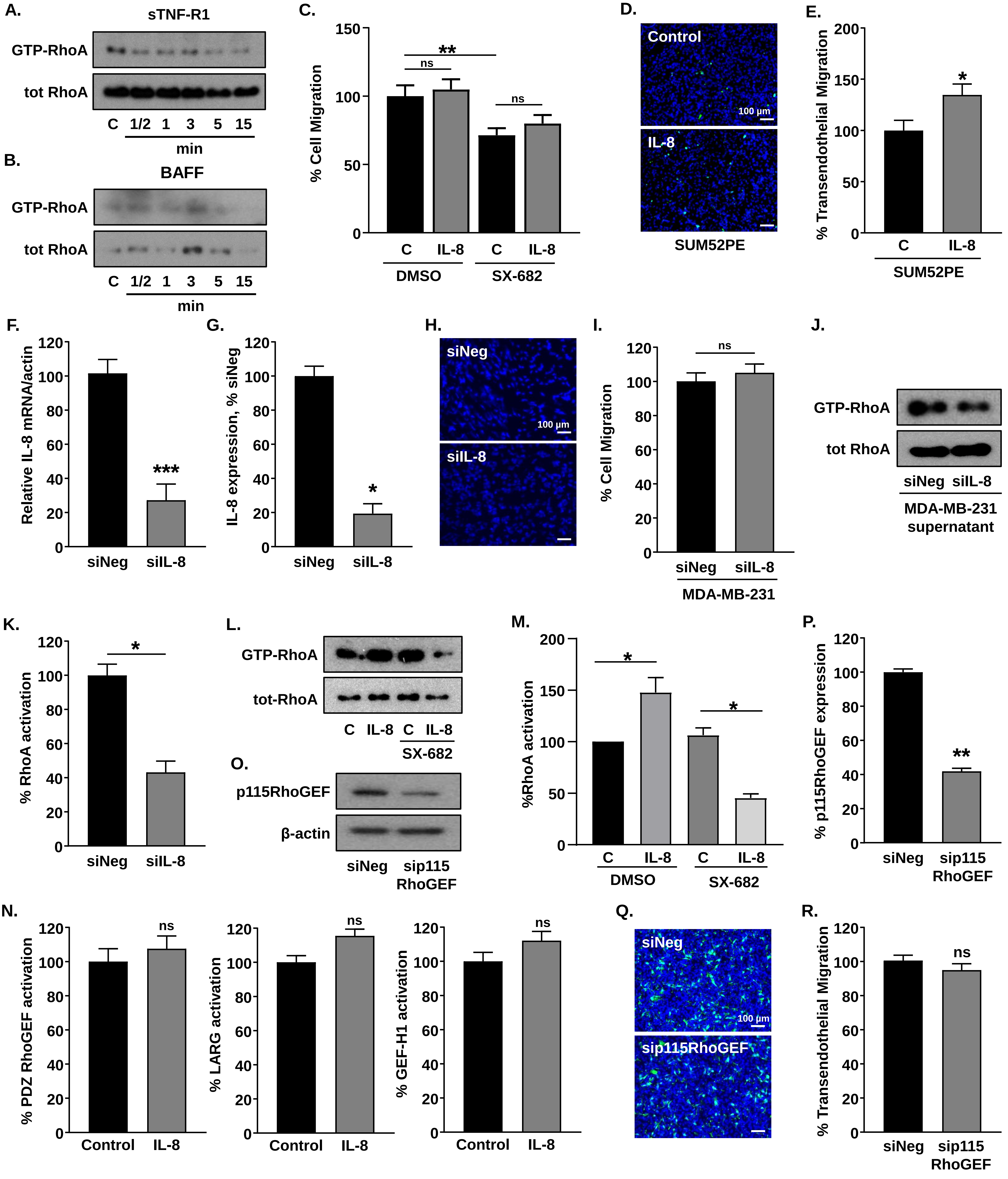

### Supplemental Figure 3.

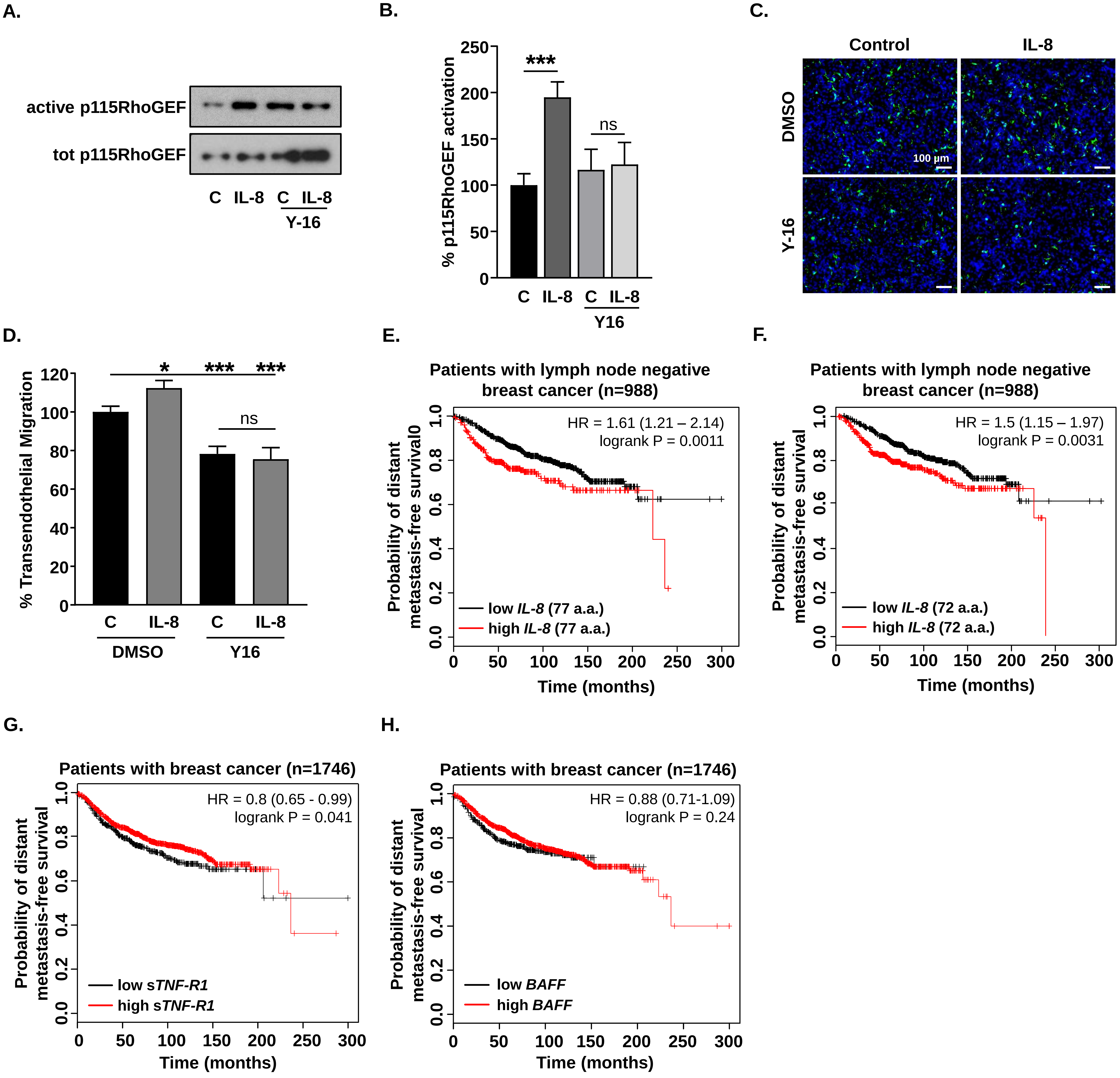

#### Supplemental Figure 4.

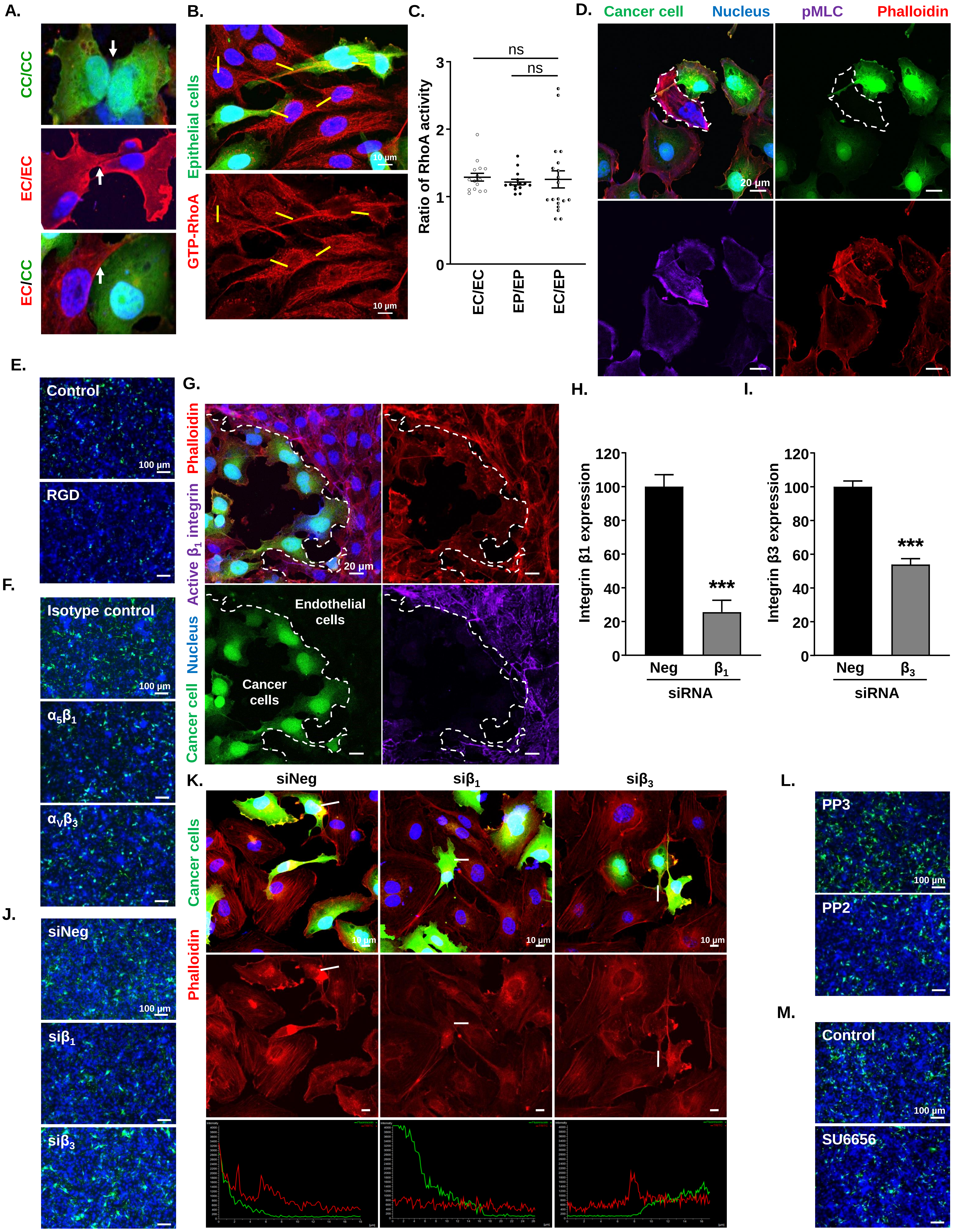

**Supplemental Figure 5.**

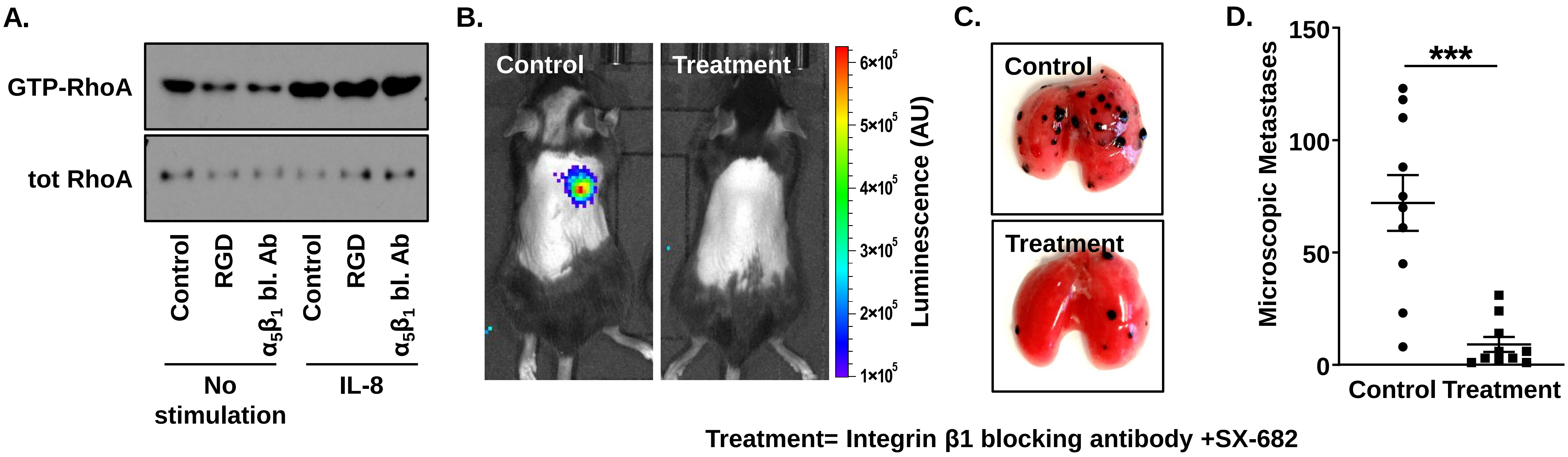


Treatment= Integrin β1 blocking antibody +SX-682

Supplemental Figure 6.

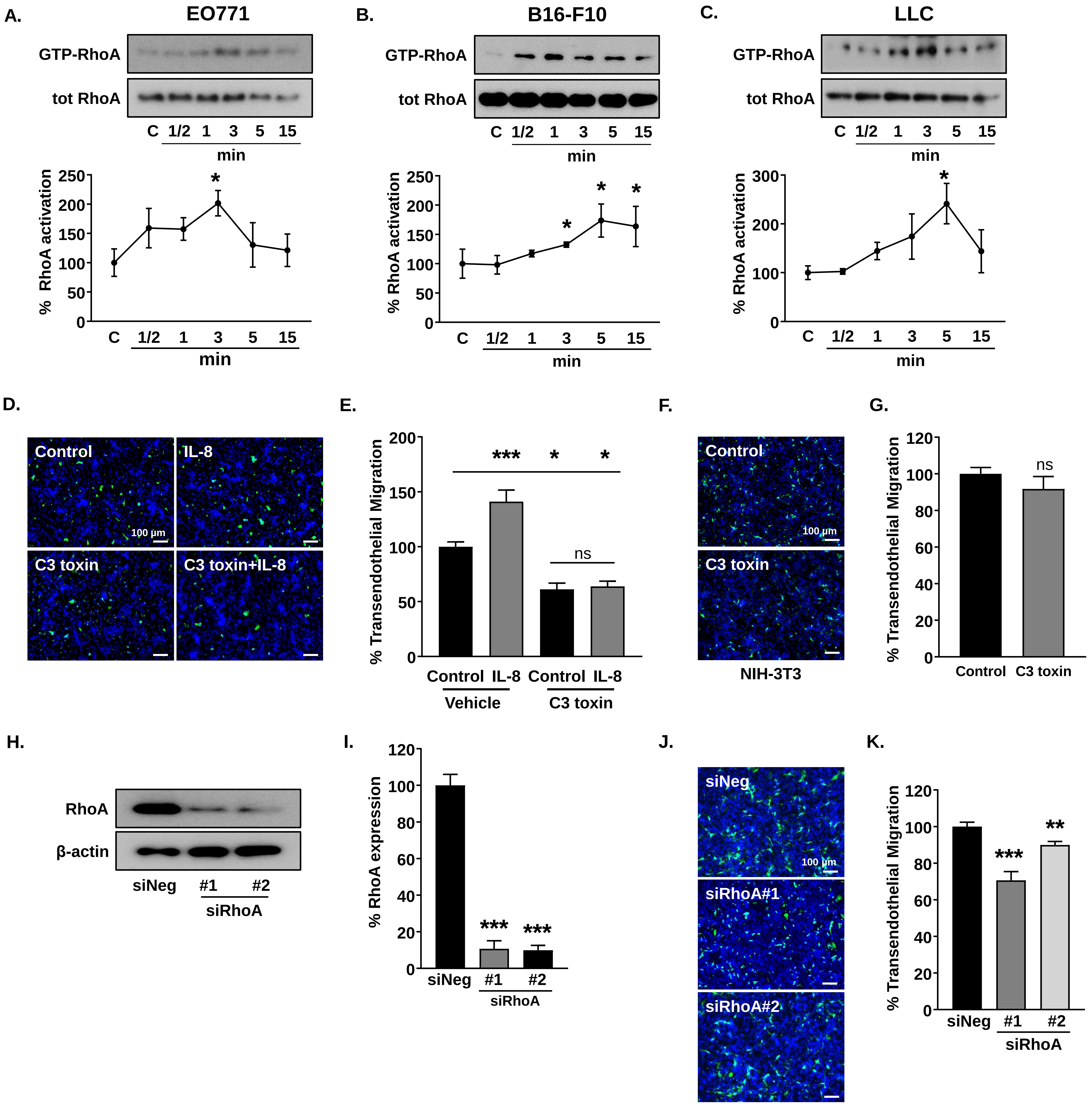

**Supplemental Figure 7.**

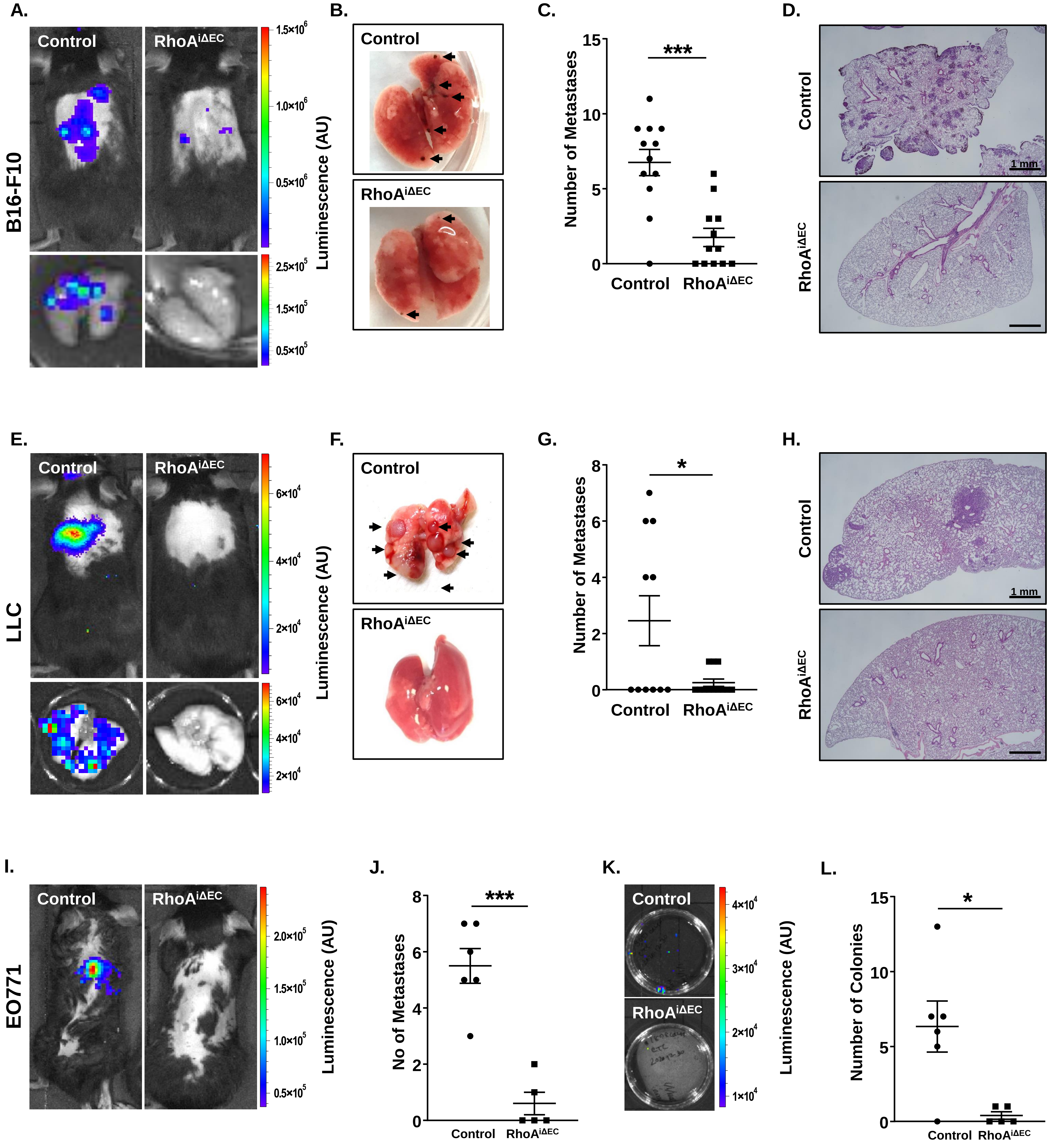

**Supplemental Figure 8.**

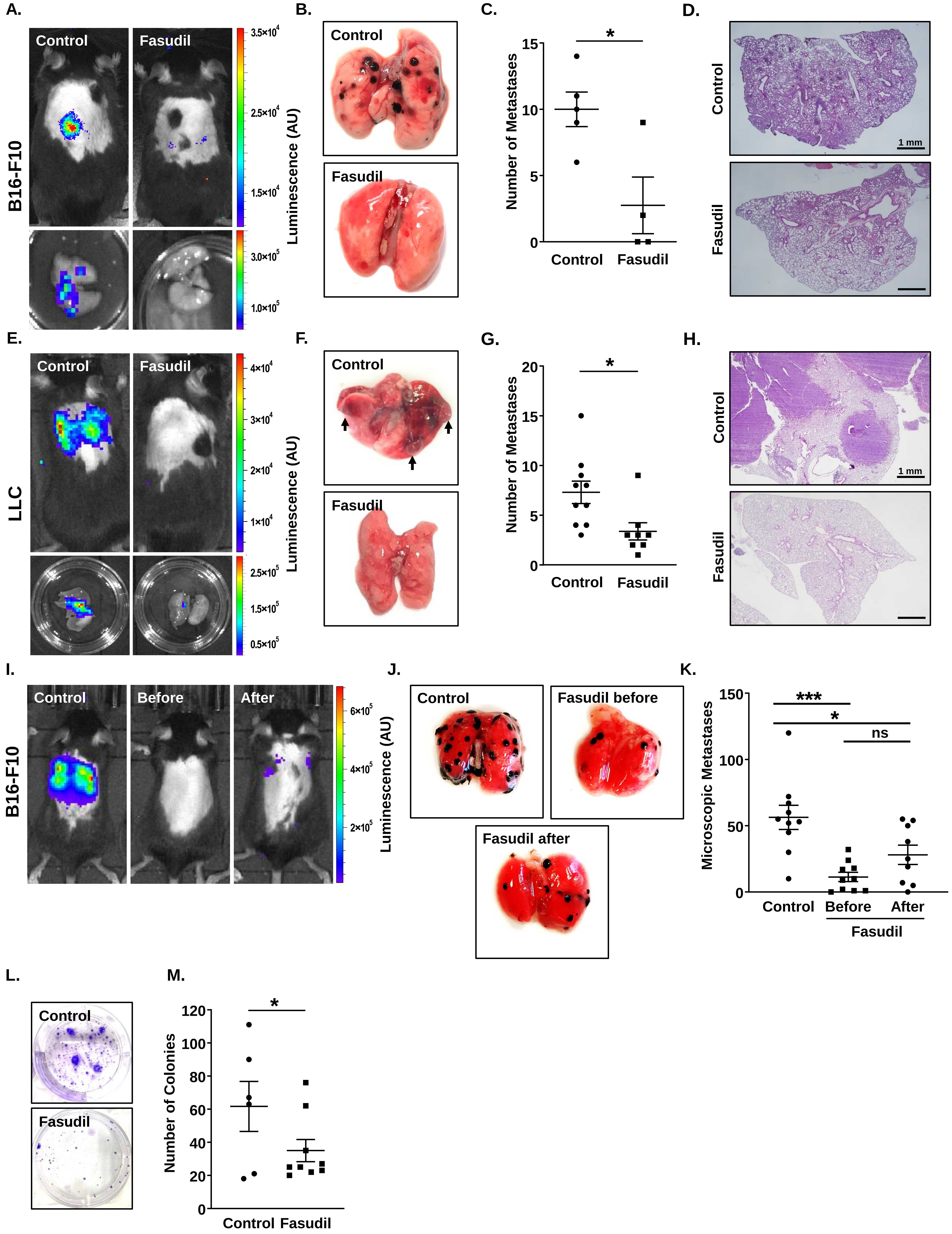

Supplemental Figure 9.

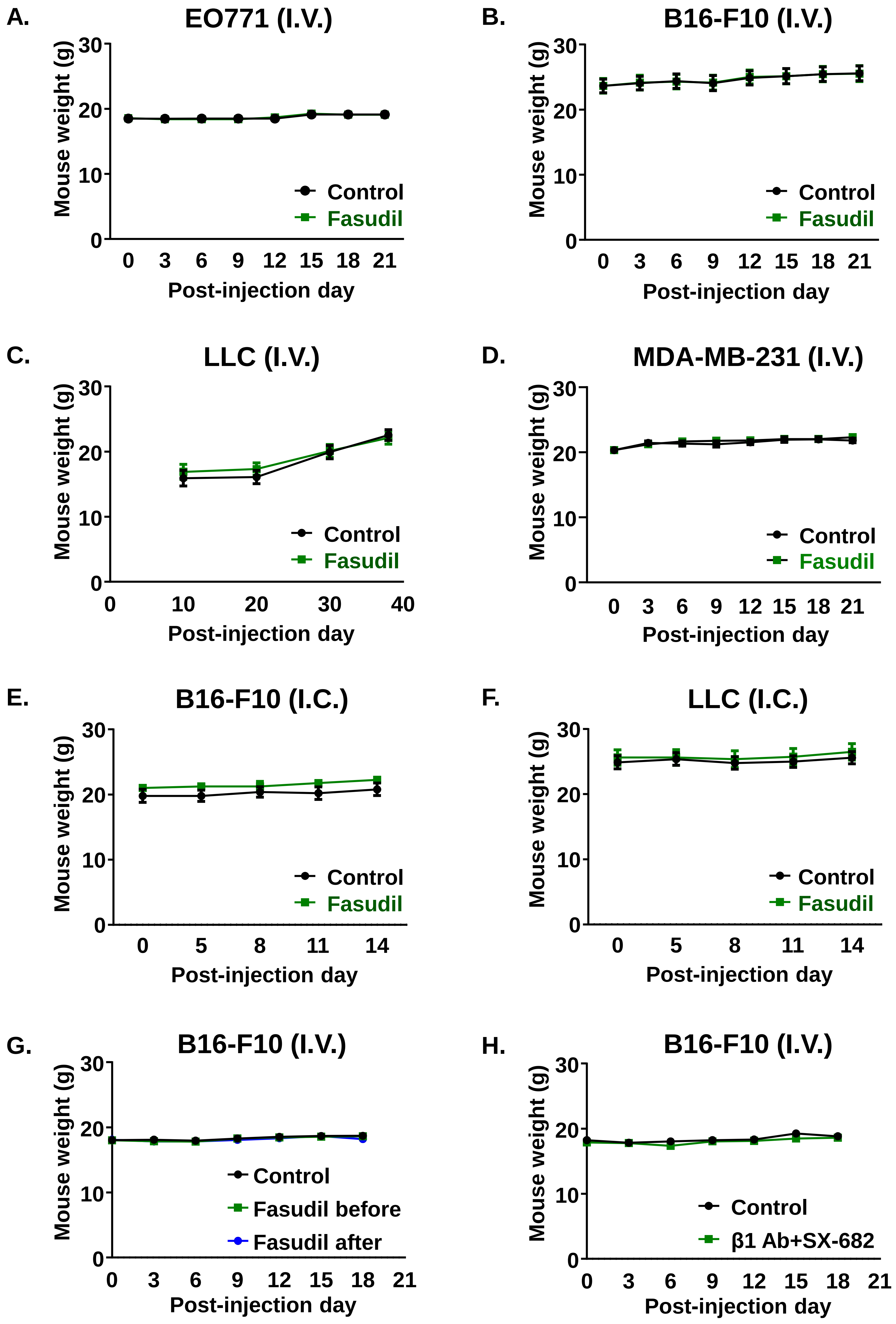
